# Bombesin-like peptide recruits disinhibitory cortical circuits and enhances fear memories

**DOI:** 10.1101/2020.10.26.355123

**Authors:** Sarah Melzer, Elena Newmark, Grace Or Mizuno, Minsuk Hyun, Adrienne C. Philson, Eleonora Quiroli, Beatrice Righetti, Malika R. Gregory, Kee Wui Huang, James Levasseur, Lin Tian, Bernardo L. Sabatini

**Affiliations:** Dept. of Neurobiology, Howard Hughes Medical Institute, Harvard Medical School, 220 Longwood Ave, Boston, Massachusetts 02115, USA; Departments of Biochemistry and Molecular Medicine, School of Medicine, University of California, Davis, Davis, CA, USA

**Keywords:** Gastrin-releasing peptide, neuropeptide, cortex, fear memory, VIP cells, disinhibition, CRISPR/Cas9

## Abstract

Disinhibitory neurons throughout the mammalian cortex are powerful enhancers of circuit excitability and plasticity. The differential expression of neuropeptide receptors in disinhibitory, inhibitory and excitatory neurons suggests that each circuit motif is controlled by distinct neuropeptidergic systems. Here, we reveal that a bombesin-like neuropeptide, gastrin-releasing peptide (GRP), recruits disinhibitory cortical microcircuits through selective targeting and activation of vasoactive intestinal peptide (VIP)-expressing cells. Using a newly-developed genetically-encoded GRP sensor and trans-synaptic tracing we reveal that GRP regulates VIP cells via extrasynaptic diffusion from several putative local and long-range sources. *In vivo* photometry and CRISPR/Cas9-mediated knockout of the GRP receptor (GRPR) in auditory cortex indicate that VIP cells are strongly recruited by novel sounds and aversive shocks, and that GRP-GRPR signaling enhances auditory fear memories. Our data establish peptidergic recruitment of selective disinhibitory cortical microcircuits as a mechanism to regulate fear memories.

## Introduction

Cortical circuits consist of multiple cell classes whose orchestrated activity is crucial for signal processing and plasticity. The post-synaptic specificity of afferent and intracortical inputs permits the temporally precise regulation of different cortical cell classes during behavior (Karnani et al., 2016; Pfeffer et al., 2013; Pi et al., 2013). Distinct thalamo-cortical inputs convey sensory information through feed-forward excitatory circuits (Bruno and Sakmann, 2006), whereas cortico-cortical synaptic inputs mediate top-down or cross-modal control of sensory processing dependent on predictions about the environment, attentional state, emotional valence and behavioral context (Buschman and Miller, 2007; Desimone and Duncan, 1995; Fontanini and Katz, 2006; Gregoriou et al., 2009; Iurilli et al., 2012; Lee et al., 2013; Moore and Armstrong, 2003; Saalmann et al., 2012; Sundberg et al., 2009). For example, cortical inputs facilitate mismatch signals (Leinweber et al., 2017), suppress responses to predicted and unattended stimuli (Iurilli et al., 2012), or enhance sensory responses and plasticity (Fu et al., 2015; Lee et al., 2013; Zhang et al., 2014) by targeting mainly excitatory, inhibitory or disinhibitory cortical neurons, respectively. Thus, cortical circuit motifs, defined by stereotyped synaptic connectivity between transcriptionally- and functionally-distinct cell classes, are regulatory control points that can be differentially activated to induce behaviorally-relevant changes in cortical state.

The specificity of the expression in cortical neurons of receptors for neuromodulators, signaling molecules that often act through slower extrasynaptic transmission, suggests that, in parallel to these fast-acting inputs, multiple channels of cortical neuromodulator/neuropeptide-based communication exist that also regulate functionally-relevant cellular and network activity (Smith et al., 2019; Tasic et al., 2016). For example, the neuropeptide oxytocin controls cortical signal processing and behavior through cell type-specific modulation of inhibition, and its context-dependent release induces network state changes that support maternal and sociosexual behaviors (Marlin et al., 2015; Nakajima et al., 2014). Separately, the cellular and network effects of the neuromodulators norepinephrine (NE) and acetylcholine (ACh) induce states of high arousal or saliency and promote experience-dependent plasticity and memory (Bear and Singer, 1986; Clark and Noudoost, 2014; Hasselmo, 2006; Kimura, 2000; Kuchibhotla et al., 2017; Polack et al., 2013). Thus, neuromodulators are often released to indicate changes in physiological and emotional states and, in turn, induce appropriate changes in cortical networks. Nevertheless, most cortical neuromodulators and neuropeptides have not been investigated in detail with respect to their cellular and behavioral effects.

Within cortex, vasoactive-intestinal peptide (VIP)-expressing neurons, due to their synaptic targets, are well-positioned to control circuit excitability and plasticity (Adler et al., 2019; Batista-Brito et al., 2017; Fu et al., 2014, 2015; Karnani et al., 2016; Pfeffer et al., 2013; Pi et al., 2013). VIP cells act mainly through the release of GABA to suppress somatostatin-expressing cells whose activity inhibits distal pyramidal cell dendrites and limits synaptic plasticity (Chen et al., 2015; Karnani et al., 2016; Pfeffer et al., 2013; Pi et al., 2013). Depending on the cortical region, VIP cells can also inhibit fast-spiking parvalbumin-expressing interneurons that powerfully regulate pyramidal cell firing (Karnani et al., 2016). Lastly, many VIP cells also release ACh and VIP, which have complex effects on neuronal and non-neuronal cell types (Von Engelhardt et al., 2007; Granger et al., 2020; Obermayer et al., 2019), and mediate cerebral vasodilation (Suzuki et al., 1984; Yaksh et al., 1987). Therefore, VIP cell activation disinhibits pyramidal cells (Karnani et al., 2016; Pfeffer et al., 2013; Pi et al., 2013), facilitates synaptic plasticity (Fu et al., 2015), and increases cortical blood flow (Suzuki et al., 1984; Yaksh et al., 1987), processes that are associated with cortical engagement, learning and memory (Adler et al., 2019; Chen et al., 2015).

VIP cells are innervated by cortical and thalamic neurons synaptically (Wall et al., 2016), but also express a diverse set of neuromodulator receptors (Smith et al., 2019; Tasic et al., 2016, 2018), making them putative targets for additional local and long-range neuromodulatory communication channels.

One neuromodulator with unknown function in most cortical brain areas is gastrin-releasing peptide (GRP), a 27 amino acid long bombesin-like peptide, that is synthetized in several brain areas in mice, rats and cats (Marcos et al., 1994; Shumyatsky et al., 2002; Wada et al., 1990) and binds to the G protein-coupled GRP receptor (GRPR) with high affinity and selectivity (Kroog et al., 1995). GRP release in different parts of the central nervous system mediates itch sensation (Sun and Chen, 2007) and sighing (Li et al., 2016b), and it has been implicated in fear memories via potential actions in the amygdala, hippocampus and prefrontal cortex, although the direction of this effect is controversial (Mountney et al., 2006, 2008; Roesler et al., 2003). Moreover, the function of GRP in most cortical areas including primary sensory and motor areas and the mechanisms by which it regulates cortical function are unknown.

Here we identify GRPR as a previously-unknown regulator of VIP cell-dependent signaling and behavior. We demonstrate that GRPR is expressed nearly exclusively in VIP cells in many cortical regions whereas its ligand, GRP, is produced by multiple long-range projecting neurons in brain regions involved in auditory fear conditioning, including amygdala and thalamus, suggesting an intra-cortical and sub-cortical mechanism of activating cortical VIP cells to induce context-dependent cortical state changes. Consistent with this hypothesis, GRP powerfully depolarizes and activates VIP cells, thereby disinhibiting pyramidal cells through direct inhibition of PV and SST neurons. These effects are sufficient to induce cell type-specific changes in immediate early gene expression. Expression of GRP by projection neurons that converge in auditory cortex but do not synapse directly onto VIP cells suggests that GRP signals via extra-synaptic diffusion rather than point-to-point transmission, consistent with results using a novel GRPR-based optical sensor that shows that GRP can diffuse in a functional form through cortical tissue. Furthermore, CRIPSR/Cas9-mediated and conditional KO of GRPR in the auditory cortex diminishes fear memories in a discriminatory auditory fear conditioning task (Letzkus et al., 2011) that we find engages VIP cells in a cue- and novelty-dependent manner. Our results thus highlight the importance of neuropeptidergic cell type-specific communication channels in regulating synaptic transmission and plasticity in functionally relevant cortical circuits.

## Results

### Cortex-wide cell type-specific expression of GRP and its receptor in mice and humans

To identify candidate neuromodulator receptors for selective regulation of VIP cells, we analyzed gene expression in mouse visual cortex in two single-cell RNA sequencing datasets (Tasic et al., 2016, 2018). Out of 11 identified genes whose expression was found to be correlated with that of *Vip* (correlation coefficient >0.5) in both datasets, the gastrin-releasing peptide receptor (*Grpr,* Fig. S1A) is the only neuropeptide receptor, and thus is a candidate for VIP cell-specific peptidergic neuromodulation (as suggested previously (Smith et al., 2019)). Fluorescent *in situ* hybridization (FISH) targeting the gene encoding its specific ligand, gastrin-releasing peptide (*Grp*), revealed strong mRNA expression across multiple cortical areas (Fig. 1A-C and Fig. S1B-C), with enrichment in L2/3 and L6 in all examined primary sensory cortical areas (Fig. 1C). Furthermore, in all analyzed areas, *Grpr^+^* cells were enriched in superficial layers, resembling the distribution of *Vip^+^* cells across cortical layers (Fig. 1D). Thus, receptor and ligand expression patterns indicate the potential for local intra-cortical GRP-GRPR signaling.

**Figure 1:**
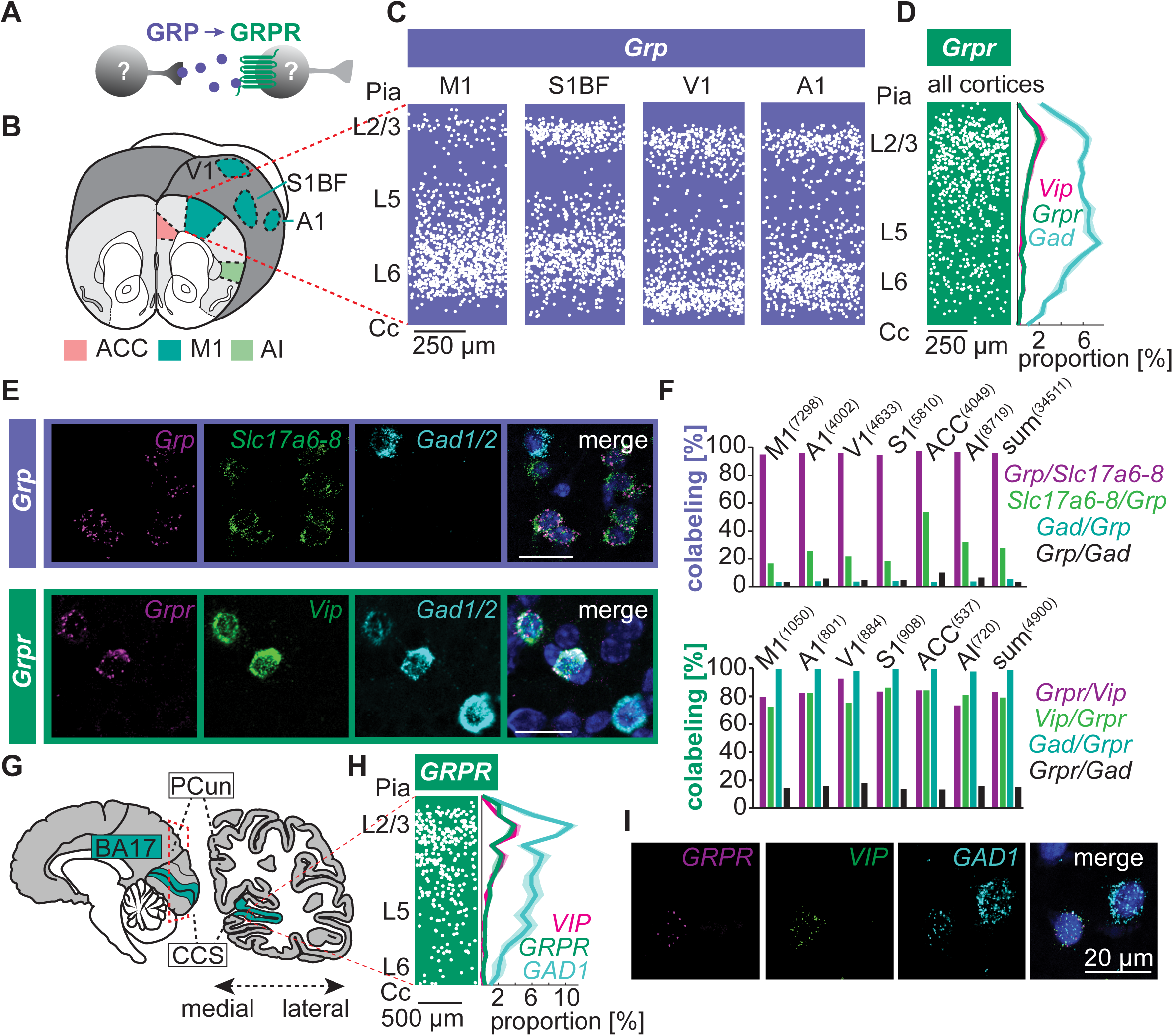
**Cortex-wide cell type-specific expression of GRP and its receptor in mice and humans.** **A**, Schematic drawing of GRP release and binding to GRP receptor (GRPR) in unidentified cell types. **B**, GRP and GRPR expression were analyzed in mice in the indicated cortical areas using fluorescent *in situ* hybridization (FISH) and confocal imaging. Abbreviations of cortical regions: M1: primary motor; A1: primary auditory; V1: primary visual; S1BF: primary somatosensory – barrel field; ACC: anterior cingulate; AI: anterior insular. **C**, Overlay showing the locations of all identified *Grp^+^* cells in the indicated cortical areas (n=5 slices for S1BF, V1, and A1 and 7 for M1). For ACC and AI summary data, see Figure S1. **D**, Overlay showing the locations of all identified *Grpr^+^* cells in the indicated cortical areas (n=34 slices, 4900 cells). Quantification of the proportions (mean ± SEM) of cells that are *Grpr^+^*, *Vip^+^* or *Gad^+^* across cortical depth (20 bins). **E**, Representative confocal images of mouse cortex showing co-expression of *Grp* (*top*) and *Grpr* (*bottom*) with *Vip*, glutamatergic markers (*Slc17a6-8* encoding vGluT1-3) and GABAergic markers (*Gad1,2*). Scale bars, 20 µm. **F**, Quantification of *Grp* (*top*) and *Grpr* (*bottom*) coexpression with indicated genes from FISH analysis as in panel E. The numbers of counted cells per brain area are indicated above bars (n=4-7 hemispheres per cortical area). **G**, Schematic showing location of human visual cortex (BA17) in which *GRPR* expression was analyzed using FISH. **H**, *Left,* Overlay showing locations of all identified *GRPR^+^* cells in 5 sections of human visual cortex. *Right,* Quantification of the proportions (mean ± SEM) of cells that are *GRPR^+^, VIP^+^* and *GAD1^+^* (n=882 cells across 20 bins). **I**, Representative confocal image of a *GRPR^+^/VIP^+^/GAD1^+^* human cell. See also Figure S1.

The layer-specific *Grp* expression resembles the expression patterns of subsets of glutamatergic cells whereas that of *Grpr* suggests, as expected, expression in *Vip* cells. Indeed, FISH analysis revealed *Grp* expression in a subset of glutamatergic cells (95.8 ± 0.4% of *Grp^+^* cells were *Slc17a6/7/8^+^*, encoding glutamatergic markers vGluT2/1/3) constituting 27.6 ± 4.9% of all glutamatergic cells across cortex (n=29841 cells; ≥ 15 slices from 3 mice per area, Fig. 1E-F). In contrast, we detected *Grp* expression in only a small subset of GABAergic cells (3.1 ± 0.1% of *Grp^+^* cells expressed *Gad1/2,* encoding GAD67/65, constituting 5.3 ± 0.9% of *Gad1/2^+^* cells; n=12868 cells; ≥ 15 slices from 3 mice per area; Fig. 1E-F). To test whether GRP is a potential modulator specifically of cortex-wide VIP cell activity, we analyzed coexpression of *Grpr*, *Vip* and GABAergic markers *Gad1/2* across multiple cortical areas in adult mice (Fig. 1B). Expression levels of *Vip* and *Grpr* were highly correlated across all areas (correlation coefficient = 0.65, Fig. S1D). *Grpr* expression was detected in 80.9 ± 2.2% of *Vip*^+^ cells, and 83.2 ± 2.6% of *Grpr^+^* cells expressed *Vip* (Fig. 1E-F). Similar numbers were found in the auditory cortex during early postnatal development (postnatal day 18, Fig. S1E) suggesting preserved GRPR-mediated signaling throughout development and adulthood.

Since 19.1% of *Grpr^+^* cells were *Vip*-negative GABAergic neurons (99.5 ± 0.3% of *Grpr^+^* cells expressed *Gad1/2*; n=4900 cells in 7 cortical areas, 4-7 hemispheres each, Fig. 1E-F), we examined whether *Grpr* is present in other major inhibitory cell types marked by *Pvalb* and *Sst* expression. Indeed, we detected *Sst* expression in 17.2% of *Grpr^+^* cells in the auditory cortex (ACx) (constituting 4.7% of *Sst^+^* cells, n=5 slices from 3 mice, Fig. S1F), whereas only 2.7% of *Grpr^+^* cells expressed *Pvalb* (0.7% of *Pvalb^+^* cells, n=5 slices from 3 mice, Fig. S1F). In contrast, in the adult mouse hippocampus, an area with comparably high expression levels of *Grpr,* expression was relatively evenly distributed among the three major GABAergic cell types identified by *Vip*, *Sst* and *Pvalb* (Fig. S1G), suggesting that preferential VIP cell targeting of GRP is a unique feature of neocortical circuits.

Several neuropeptidergic systems are highly evolutionarily conserved (Jékely et al., 2018), raising the question as to whether GRPR signaling follows similar principles in the human cortex as in mouse. Indeed, FISH analysis revealed that *GRPR^+^* and *VIP^+^* cells in the human visual cortex (BA17) follow a similar distribution across cortical layers (Fig. 1G-H) and *GRPR* mRNA was detected almost exclusively in GABAergic cells (98.2% of *GRPR^+^* neurons co-expressed *GAD1*; n=1929 cells in 12 slices; Fig. 1I and S1H). Moreover, we found that a large proportion of *VIP^+^* cells expressed *GRPR* (60.8%), and the majority of *GRPR^+^* cells expressed *VIP* (51.1%, 5 slices; Fig. 1I and S1H). In contrast, we detected *SST* and *PVALB* expression only in 1.0% and 5.6% of *GRPR^+^* cells, respectively (n=254 cells in 3 slices (*SST*) and 264 cells in 4 slices (*PVALB*); Fig. S1H). A second, non-overlapping *GRPR* oligoprobe confirmed these results (Fig. S1H), suggesting that the neocortical GRP-GRPR signaling pathway is evolutionarily conserved between mouse and human. The large fraction of human *GRPR^+^* cells that lack *VIP* expression may represent a distinct *VIP/SST/PVALB*-negative cell class or cells in which *VIP* mRNA levels are below detection threshold.

### Putative local and long-range sources of synaptic and extrasynaptic GRP

The strong expression of *Grp* in glutamatergic cells in L6 of all examined primary sensory and motor cortical areas raised the question as to whether *Grp* is selectively expressed in one of the two major glutamatergic L6 neuronal cell types: cortico-cortical and cortico-thalamic projection neurons (Briggs and Usrey, 2011). To identify cortico-thalamic neurons, we injected Cholera Toxin B (CTB) into the auditory thalamus (Fig. 2A-B) resulting in retrograde labeling of L6 cortico-thalamic neurons in the auditory cortex (Fig. 2B-C). FISH revealed that 86.3 ± 4.0% of *Grp^+^* cells in L6 were retrogradely labeled with CTB (n=958 *Grp^+^* cells counted in 4 hemispheres) and that L6 *Grp*^+^ neurons constitute a subpopulation of cortico-thalamic neurons (*Grp* was expressed in 40.7 ± 9.9% of CTB labeled cells in L6; n=1494 CTB^+^ cells counted in L6 in 4 hemispheres; Fig. 2C). Similarly, injections into the motor thalamus (mainly ventral lateral and posterior nuclei) resulted in retrograde labeling of 72.0 ± 7.1% of *Grp*^+^ L6 neurons in motor cortex and 55.3 ± 5.9% of CTB^+^ cells in L6 were *Grp*^+^ (n=1585 CTB^+^ cells and 975 *Grp^+^* cells counted in L6 of the motor cortex in 3 hemispheres), establishing L6 cortico-thalamic projection neurons as a putative source of cortical GRP signaling.

**Figure 2:**
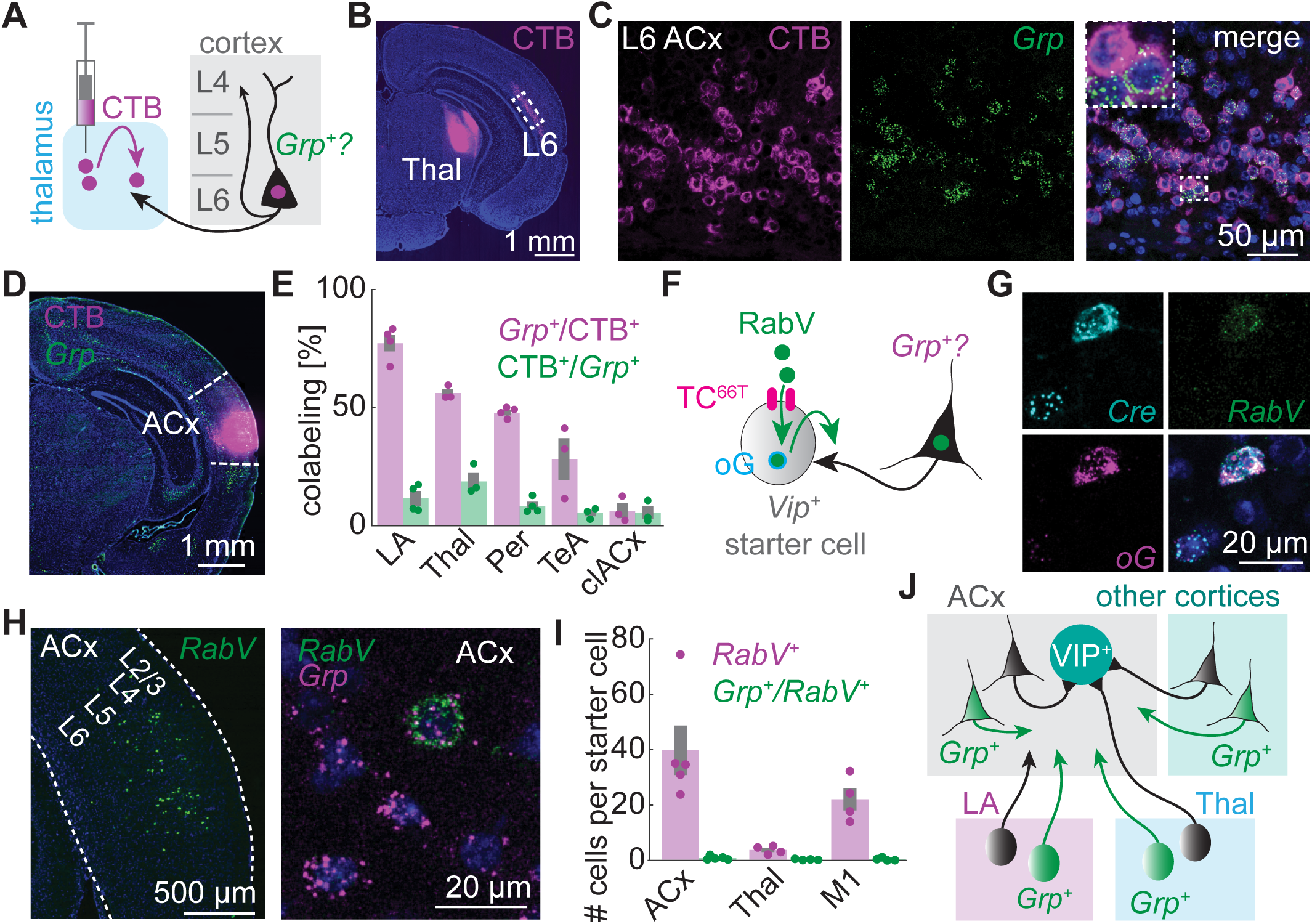
**Putative local and long-range sources of synaptic and extrasynaptic GRP** **A**, Schematic of retrograde tracing with CTB injection into auditory thalamus to quantify *Grp* expression in L6 cortico-thalamic neurons. **B**, Representative epifluorescent image showing CTB injection into auditory thalamus and retrograde labeling in auditory cortex L6. **C**, Confocal image of retrogradely labeled cells in layer 6 of auditory cortex and FISH against *Grp*. Inset: magnification of the highlighted area. **D**, Retrograde tracing with CTB injected into auditory cortex to quantify *Grp* expression in corticopetal projection neurons. Epifluorescent image of a representative CTB injection and FISH against *Grp*. **E**, Quantification of *Grp^+^* cells among CTB^+^ retrogradely labeled cells (magenta) and vice versa (green) in the indicated areas following CTB injection as in panel D. Each dot represents data from one mouse. Mean ± SEM across 3-4 mice per area. **F**, Schematic drawing of the components used for transsynaptic tracing from *Vip*^+^ starter cells using pseudotyped rabies virus SADΔG-EnVA-H2B-EGFP (RabV). **G**, Confocal image of an exemplary starter cell in auditory cortex identified by *RabV-gp1* (targeting the RabV nucleoprotein N), *oG* and *Vip^Cre^* coexpression (FISH). **H**, Confocal images of FISH analysis showing *RabV-gp1^+^* cells in auditory cortex (*left*) and of an exemplary *Grp^+^* retrogradely labeled (*RabV-gp1^+^*) cell (*right*). **I**, Quantification of numbers of retrogradely labeled cells and of those expressing *Grp* in auditory cortex (ACx) and thalamus (Thal) after injections into auditory or in motor cortex (M1) after injections into M1. Numbers of *RabV-gp1^+^* cells were normalized to the numbers of starter cells. Each dot represents data from one mouse. Mean ± SEM from n=4-5 mice. **J**, Schematic drawing of putative synaptic and extrasynaptic *Grp*-expressing inputs to auditory cortex VIP cells. See also Figure S2.

We also detected strong *Grp* expression in several input areas of the auditory cortex including the lateral amygdala (LA), posterior auditory thalamus (MGM, SG and PIL), contralateral ACx (clACx,) temporal association area (TeA) and the perirhinal cortex, suggesting that long-range projecting neurons might also be sources of cortical GRP signaling. To examine this possibility, we injected CTB into the ACx (Fig. 2D) and analyzed *Grp* expression by FISH. Indeed, the majority of retrogradely labeled cells in the LA expressed *Grp* (77.5 ± 3.5% of CTB^+^ cells; n=358 CTB^+^ cells in 15 slices from 4 mice, Fig. S2.1A). Moreover, about half of the retrogradely labeled cells in the posterior (but not anterior) thalamic suprageniculate (SG) and medial geniculate (MGM) nuclei (56.2 ± 1.7%; n=366 CTB^+^ cells in 9 slices from 3 mice; Fig. S2.1B) and in the perirhinal cortex (47.9 ± 1.1%; n=544 CTB^+^ cells in 12 slices from 4 mice) were *Grp^+^* (Fig. 2E). Less coexpression was found in the TeA (28.5 ± 8.9%; n=528 CTB^+^ cells in 9 slices from 3 mice; Fig. 2E) and clACx (6.5 ± 3.1%; n=1142 CTB+ cells in 11 slices from 3 mice; Fig. 2E). In each area, only a small population of *Grp^+^* cells were retrogradely labeled (Fig. 2E), suggesting that a subset of *Grp^+^* cells mainly in the LA and posterior auditory thalamus are putative long-range sources of GRP in cortex.

While fast synaptic neurotransmission in neuronal networks typically occurs between specific subsets of directly connected neurons, extrasynaptic diffusion of neuropeptides may allow these neuromodulators to reach all cells in a target region. To test whether VIP cells receive synaptic inputs and peptidergic signals from overlapping or distinct subsets of neurons, we used pseudotyped rabies virus (RabV) transsynaptic retrograde labeling to identify cells that provide synaptic inputs to VIP cells. To restrict transsynaptic retrograde labeling to VIP cells, we injected helper viruses encoding Cre-dependent avian receptor TVA with increased specificity (AAV2/9 CAG-DIO-TC^66T^-mCherry (Miyamichi et al., 2013)) and Cre-dependent optimized glycoprotein (AAV2/9 CAG-DIO-oG (Kim et al., 2016)) into the motor and auditory cortices of *Vip-IRES-Cre* mice. Three weeks later we injected glycoprotein-deleted pseudotyped rabies virus encoding nuclear localized EGFP (SADΔG-H2B:EGFP(EnvA) (Mandelbaum et al., 2019)) into the same region (Fig. 2F). *Vip^+^* starter cells (*Vip-Cre^+^* cells expressing *oG* and RabV detected by FISH using the *RabV-gp1* probe targeting RabV nucleoprotein N, Fig. 2G) in the ACx were detected over a range of 400-720 μm (anterior-posterior axis, n=5 mice). Retrogradely labeled cells were located in the ipsilateral (Fig. 2H) and contralateral ACx, auditory thalamus and ipsilateral higher order auditory and association cortices, but not in the LA, consistent with previous reports (Wall et al., 2016). Importantly, AAV2/9 CAG-DIO-TC^66T^-mCherry injection did not lead to significant RabV labeling in the absence of Cre (3 RabV^+^ cells across 10 injections into auditory and motor cortices, CTB was coinjected with RabV in 2 cases to verify successful injections; Fig. S2.1C), confirming that helper virus expression and RabV entry were highly-specific to Cre-expressing VIP cells. Moreover, AAV2/9 CAG-DIO-oG injection into wild type mice did not lead to any detectable transsynaptic RabV labeling in the absence of Cre even when RabV infectivity was permitted at high levels using a Cre-independent version of TC^66T^-mCherry (0 RabV^+^/TC^66T-^ cells among 5740 RabV^+^/TC^66T+^ cells, 2 injections into motor cortex and 2 into ACx; Fig. S2.1D).

To test whether *Grp* is expressed in neurons providing synaptic inputs to VIP cells, we detected neurons infected with RabV by FISH using the *RabV-gp1* probe (Fig. 2H), and we analyzed *Grp* expression in retrogradely-labeled *RabV-gp1^+^* cells. Only a small subpopulation of *RabV-gp1^+^* neurons expressed *Grp* in the cortex (2.1 ± 0.4%; 0.9 ± 0.3 *Grp^+^/ RabV-gp1^+^* cells per starter cell, 39.8 ± 8.9 *RabV-gp1^+^* cells per starter cell, n=1348 retrogradely labeled cells counted in ACx of 5 mice, Fig. 2H-I), and in the posterior thalamus (7.9 ± 1.9%; 3.8 ± 0.7 *RabV-gp1*^+^ cells per starter cell, 0.3 ± 0.1 *Grp^+^/ RabV-gp1^+^* cells per starter, Fig. 2I and Fig. S2.1E). Similarly, low numbers of *RabV-gp1^+^/Grp^+^* cells were detected after injections of RabV and helper viruses into the M1 of *Vip-IRES-Cre* mice (Fig. 2I and Fig. S2.1F), suggesting that VIP cells across multiple cortical areas receive GRP signals from a neuronal population that is largely distinct from that which provides synaptic input. Thus, GRP-GRPR signaling is a previously unknown pathway for extrasynaptic cell-to-cell communication to cortical VIP cells (Fig. 2J).

Extrasynaptic GRP signaling requires that the peptide be stable and efficiently diffuse through extracellular space. To image GRP diffusion *in vivo*, we developed a genetically-encoded GRP sensor based on a previously established platform (Fig. S2.2A-B (Patriarchi et al., 2018)), by replacing intracellular loop 3 of human GRPR with circularly permuted GFP (cpGFP). The dynamic range and affinity of the resulting GRP sensor (grpLight) were further optimized by changing the compositions of the linker between cpGFP and GRPR and of the intracellular loop (Fig. S2.2C). Multiple versions of grpLight showed high specificity compared to other common neuropeptides (Fig S2.2D) and could detect nanomolar concentrations of GRP when expressed in HEK 293 cells, cultured neurons from rat hippocampi and acute brain slices (Fig. S2.2E-G). *In vivo* expression and dual-color photometric imaging (Fig S2.2H) of grpLight in the auditory cortex following infusion of red fluorescently-tagged GRP (TAMRA-GRP) into a distant visual cortical area (Fig S2.2I) revealed long-lasting fluorescence increases minutes after the start of GRP infusion (Fig S2.2J). Our results show that GRP diffuses slowly and maintains biological activity for over an hour in intact brain tissue, suggesting that GRP is a long-acting peptide that likely arises from non-synaptic sources and reaches large neuronal populations through extrasynaptic diffusion and long-range volume transmission.

### GRP depolarizes and increases intracellular calcium in cortical VIP cells

The cell type specificity of *Grpr* expression suggests that VIP cells are the primary targets of functional modulation by GRP. Whole-cell current-clamp recordings in Vip-EGFP^+^ neurons in the ACx of male mice in the presence of glutamate and GABA neurotransmission blockers (10 µM NBQX, 10 µM CPP, 10 µM gabazine, 2 µM CGP55845) (Fig. 3A and Fig. S3A) revealed that most VIP cells (7 out of 10) depolarize upon GRP bath application (300 nM for 2 min, mean depolarization: 5.8 ± 1.2 mV), partially resulting in long-lasting (>1 min) burst-like firing activity (3 out of 10 neurons, Fig. 3B and S3B). Depolarization was concentration-dependent (Fig. 3C) and strongly reduced by the specific GRPR antagonist BW2258U89 (10 µM: mean depolarization 0.70 ± 0.58 mV, t(14)=-4.28, p<0.001; Fig. S3B). As the *Grpr* gene is on the X-chromosome, we separately examined VIP cell responses in female mice and found a slightly, but not significantly, larger depolarization (8.62 ± 1.11 mV; n=10 cells; t(18)=1.95, p=0.07; Fig. S3C) compared to male mice. Similar to VIP cells in ACx, 9 out of 10 tested VIP cells in M1 depolarized in response to GRP application, and 3 out of 10 neurons developed burst-like activity (Fig. S3E).

**Figure 3:**
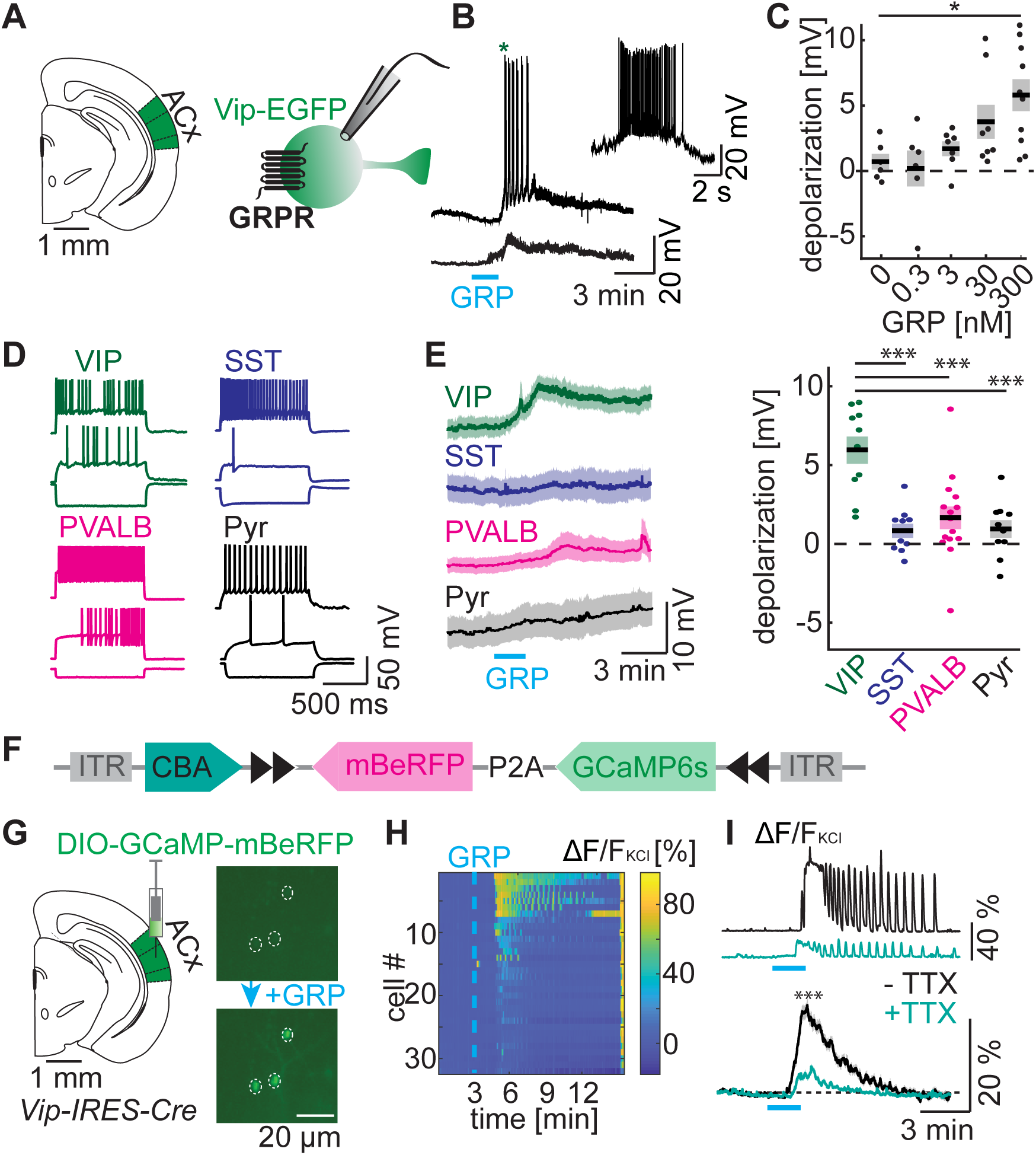
**GRP depolarizes and increases intracellular calcium in cortical VIP cells** **A**, Schematic showing whole-cell current-clamp recordings of VIP cells in auditory cortex of Vip-EGFP mice used to examine effects of GRP application. **B**, Two exemplary VIP cells responding with bursts (*top*) or depolarization (*bottom*) following 2 min bath application of GRP (300 nM) in the presence of synaptic blockers (NBQX, CPP, gabazine, CGP). Inset: magnification of the first burst in the upper trace indicated by an asterisk. **C**, Depolarization of VIP neurons upon application of indicated concentrations of GRP. Mean ± SEM across 6-10 cells per group. Comparison 0 vs. 300 nM GRP: 2-sample t-test for unequal variance: t(12.52)=3.76, p=0.01. Other comparisons n.s. **D**, Representative firing patterns of auditory cortex VIP, SST, PVALB and pyramidal (Pyr) cells as indicated upon -200 pA current injection (*bottom*), at AP threshold (*middle*), and at maximal firing frequency (*top*). **E**, Average time course (*left*) and amplitude (*right*) of the membrane potential changes in the indicated cell types in auditory cortex following GRP application. Mean ± SEM from n=10 VIP, SST, and pyramidal cells (Pyr) and n=15 PVALB cells. Bath contains NBQX, CPP, gabazine, CGP, TTX. Comparison of depolarizations across cell types: Bonferroni-corrected t-test: SST: t(18)=-5.27, p<0.001; PVALB: t(23)=- 3.83, p<0.001; pyramidal: t(18)=-4.87, p<0.001. **F**, Design of the plasmid for Cre-dependent stoichiometric expression of GCaMP and the large-stokes shift red fluorescent protein mBeRFP for imaging of calcium entry and detection of infected cells, respectively. **G**, Injection of AAV DIO-GCaMP-P2A-mBeRFP into auditory cortex of male *Vip^Cre^* mice (*left*) and epifluorescent GCaMP imaging (*right*) in acute slices before (*top*) and after (*bottom*) GRP bath application (300 nM, 2 min), in the presence of synaptic blockers NBQX, CPP, gabazine and CGP. **H**, Heatmap of fluorescence changes (expressed relative to fluorescence following KCl application) across all imaged VIP cells in an exemplary acute auditory cortex slice. **I**, GCaMP fluorescence changes (normalized to fluorescence following KCl application) with or without TTX in two exemplary VIP cells (*top*) or across all recorded cells (*bottom*). Mean ± SEM, Mann-Whitney U test: U=10178, p<0.0001, n=218 and 179 cells in 7 and 6 slices without and with TTX respectively. See also Figure S3.

To confirm that the effects of GRP are largely VIP cell-specific, we obtained current-clamp recordings from *Sst*-EGFP, *Pvalb*-EGFP cells and pyramidal cells (identified based on morphological and electrophysiological properties) in the ACx (Fig. 3D) and M1 (Fig. S3F) using synaptic blockers (NBQX, CPP, gabazine, CGP) and TTX to abolish all activity-dependent neurotransmission. GRP-evoked depolarization in all three cell types was significantly smaller than in VIP cells in the ACx (SST: 0.76 ± 0.45 mV, t(18)=-5.27, p<0.001; PVALB: 1.58 ± 0.71 mV, t(23)=-3.83, p<0.001; pyramidal: 0.87 ± 0.56 mV, t(18)=-4.87, p<0.001; VIP: 5.87 ± 0.86 mV; Fig. 3E) as well as in the M1 (SST: 1.39 ± 0.74 mV, U=2, p=0.001; PVALB: 1.78 ± 0.45 mV, U=0, p=0.001; pyramidal: 0.56 ± 0.49 mV, U=0, p=0.001; VIP: 9.38 ± 1.28 mV; Fig. S3G). No significant difference was found when comparing PVALB cell depolarization upon GRP application to control recordings without GRP (n=13 PVALB cells with 0 nM GRP; U=76, p=0.33, Fig. S3D), confirming that VIP cells are the preferential target of GRP signaling.

Previous reports suggested that GRPR is a Gα_q_-coupled receptor at least in some cell types (Hellmich 1997). A common secondary messenger of Gα_q_ signaling is intracellular calcium (Ca^2+^). To visualize Ca^2+^ dynamics in VIP cells upon GRP application, we expressed Cre-dependent GCaMP6s in *Vip-IRES-Cre* mice. To facilitate the identification of infected VIP cells in acute slices with low baseline GCaMP fluorescence, we expressed the large-stokes shift red fluorophore mBeRFP (excitation peak at 446 nm; (Yang et al., 2013)) stoichiometrically with GCaMP using a self-cleaving P2A peptide linker (Fig. 3F). Bath application of GRP increased GCaMP fluorescence in most VIP cells in the ACx of male and female mice (Median (IQR): 13.3 (18.8)% ΔF/F_KCl_ vs 11.8 (15.6)% in male vs female; U=413360, p=0.047; n=1510 and 580 cells in 37 and 14 slices respectively; Fig. 3G and Fig. S3H). Slightly larger calcium increases were found in M1 (U=233659, p<0.0001, n=405 cells in 8 motor cortex slices of male mice, Fig. S3H). Reminiscent of the GRP-induced burst activity in some VIP cells (Fig. 3B and Fig. S3E), GCaMP fluorescence exhibited non-synchronized phasic (oscillatory) fluctuations at various frequencies in 26.2% of imaged VIP cells (n=218 cells in the ACx of male mice, in the presence of NBQX, CPP, gabazine, CGP, Fig. 3H and Fig. S3I). Fluorescence increases and fluctuations were partially blocked by TTX (14.0% out of 179 cells exhibited calcium oscillations; Median (IQR) ΔF/F_KCl_: 7.43 (16.8)% without TTX vs 1.5 (2.0)% with TTX, U=10178, p<0.0001, Fig. 3H-I and Fig. 3J-K), suggesting that the majority of GRP-induced Ca^2+^ signaling is action potential-dependent. The action potential-independent Ca^2+^ elevations are consistent with regulation of intracellular Ca^2+^ release from internal stores by Gα_q_- and IP_3_-mediated signaling.

The long delay between GRP application and changes in membrane potential and intracellular Ca^2+^ may arise from delays in the infusion system (estimated at ∼1 min) and slow diffusion of the peptide through brain tissue. Indeed, separate analyses of the latency of GRP-triggered Ca^2+^ increases and of fluorescence increases following bath application of fluorescently-tagged GRP (TAMRA-GRP) revealed that Ca^2+^ increases occurred with a short latency (12 sec) after GRP levels increased (TAMRA-GRP latency: 87.4 sec (range: 79-95 sec); peak latency: 160.3 sec (range: 156.4-162.8 sec); n=3 slices; GCaMP latency: 99.4 ± 0.9 sec; peak: 156.9 ± 1.5 sec in 1372 responding VIP cells out of 1510 total cells in ACx of male mice). In summary, our data indicate that GRP is a selective modulator of VIP cell signaling, which increases the membrane potential and elevates intracellular Ca^2+^ with prolonged effects at low nM concentrations. Unfortunately, as our most sensitive grpLight sensors have apparent affinities (EC50) of ∼100-360 nM (Fig. S2.2F), they are unlikely to be able to detect physiologically-meaningful GRP levels that occur at approximately 10% of this concentration.

### GRP disinhibits cortex and induces immediate early gene expression

The main circuit effect of optogenetic VIP cell activation in the cortex is disinhibition of pyramidal cells through inhibition of SST and PVALB cells (Karnani et al., 2016; Pi et al., 2013) (Fig. 4A). To test whether the specific GRP-mediated changes in VIP cell activation and Ca^2+^ signaling lead to similar network effects, we recorded inhibitory postsynaptic currents (IPSCs) in SST and PVALB cells in acute cortical brain slices, under pharmacological block of synaptic glutamatergic neurotransmission (NBQX, CPP). Indeed, bath application of GRP increased IPSC frequencies (>2 SD above baseline) in 60% of SST cells in the ACx (mean increase: 6.08 ± 2.29 Hz; n=10 cells; Fig. 4B-C). Blocking action potentials with TTX blocked this increase (0 out of 10 SST cells responding, average increase: 0.09 ± 0.12 Hz, U=21, p=0.03). Consistent with the lower connectivity rate of VIP to PVALB cells (Karnani et al., 2016), we found increased IPSC frequencies upon GRP application in only 40% of tested PVALB cells, and the average increase was smaller than in SST cells (1.99 ± 1.05 Hz; n=10 cells; Fig. 4C). IPSC frequencies were increased only in 1 out of 10 pyramidal cells (mean increase: 0.58 ± 0.53 Hz; Fig. 4C). Similar effects were observed in M1 (increased IPSC frequency in 4 out of 10 SST cells; mean increase: 13.28 ± 5.97 Hz; 3 out of 10 PVALB cells, mean increase: 4.59 ± 2.00 Hz; Fig. S4A-B). In contrast to IPSC frequencies, EPSC frequencies were largely unaffected by GRP bath application: thus, GRP application increased EPSC frequency only in 1 out of 11 PVALB and 1 out of 10 SST cells in ACx (mean increase in EPSC frequencies: PVALB: 1.95 ± 1.43 Hz; SST: 1.86 ± 1.15 Hz), suggesting that inhibition of SST cells is the main direct network effect of GRP-mediated VIP cell activation.

**Figure 4:**
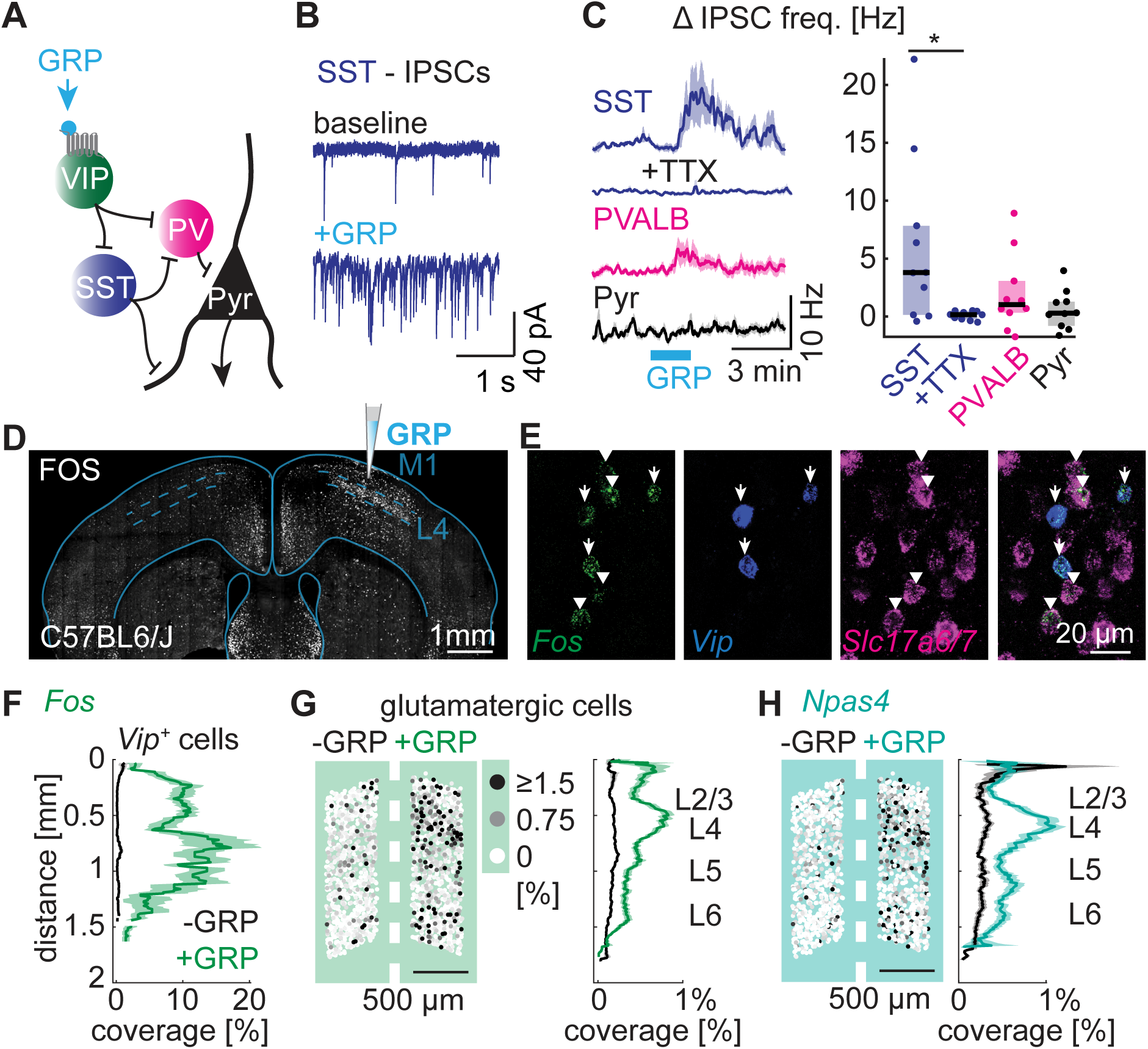
**GRP disinhibits the cortex and induces immediate early gene expression** **A**, Schematic drawing of the disinhibitory circuit that underlies VIP cell function in cortex. **B**, Whole-cell voltage-clamp recordings of IPSCs in a representative SST cell in auditory cortex before (*top*) and following (*bottom*) GRP application (300 nM, 2 min) in the presence of NBQX and CPP. **C**, Time courses (*left*, mean ± SEM) and magnitude (*right*, median and IQR) of IPSC frequency changes in SST, PVALB and pyramidal cells (n=10 cells per group). For SST cells, the effects with and without TTX present were compared (Mann-Whitney U test: U=21, p=0.03). **D,** Representative epifluorescent image of a FOS immunostaining after unilateral injection of 3 µM GRP, as schematized, into the right motor cortex in anaesthetized mice. **E**, Confocal images of *Fos* expression in *Vip^+^* (arrows) and glutamatergic cells (arrowheads) analyzed using FISH. **F-H**, *Fos* and *Npas4* expression levels (defined as cell area covered with FISH labeling) were quantified in *Vip^+^* and glutamatergic cells across all cortical layers for the right (green) and left (black) motor cortices (mean ± SEM). Intensity-coded map of expression levels (percent coverage) of all glutamatergic cells in an exemplary slice shown on the left (G, H). N=398 (*Vip*), 15108 (*Slc17a6,7*) cells, 2-3 slices from 3-5 mice for each condition and cell type. See also Figure S4.

Since VIP cell-mediated disinhibition of pyramidal cells strongly depends on the activity level of SST cells, we examined GRP-mediated disinhibition of pyramidal cells *in vivo*. To this end, we injected GRP into M1 of anaesthetized mice and used expression of the activity-induced immediate early gene *Fos* as an indicator of neuronal activity (Sheng et al., 1990). Immunostaining and FISH analysis revealed a strong increase in Fos (*Fos*) expression ipsilateral but not contralateral to the injection site (Fig. 4D and Fig. S4C). Cell type-specific comparison of *Fos* expression in both hemispheres attributed this effect to significantly increased *Fos* expression across most cortical layers in *Vip^+^* and glutamatergic cells, but not *Sst^+^* and *Pvalb^+^* cells ipsilateral to the injection site (Fig. 4E-G and Fig. S4D_1-2_). Importantly, GRP injections into M1 of mice lacking GRPR specifically in *Vip^+^* cells (*Vip-IRES-Cre;Grpr^fl/y^* (Yu et al., 2017); Fig. S4E_1-2_) or injections of the diluent (NRR) alone (Fig. S4F_1-2_) led to only small increases in *Fos* expression in both *Vip^+^* and glutamatergic cells in the injected hemisphere compared to the contralateral side. For direct statistical comparison of *Fos* expression across different conditions we plotted the cumulative distribution of expression levels in all *Vip^+^* and glutamatergic cells after normalizing *Fos* expression to the average *Fos* expression on the contralateral hemisphere (per slice and cell type; Fig. S4G) to account for movement- and staining-related differences in *Fos* labeling. Direct comparison of these data confirmed significantly smaller *Fos* expression in *Vip^+^* and glutamatergic cells when GRP was injected into *Vip-IRES-Cre;Grpr^fl/y^* mice (Kolmogorov-Smirnov: *Vip^+^* cells: p<0.0001, t=0.51; glutamatergic cells: p<0.0001, t=0.67; Fig. S4G) or when NRR was injected into control mice (Kolmogorov-Smirnov: *Vip^+^* cells: p<0.0001, t=0.87; glutamatergic cells: p<0.0001, t=0.80, Fig. S4G), indicating that GRP leads to disinhibition of glutamatergic cells through VIP cell-specific GRPR signaling.

A previous study indicated that *Fos* expression can be induced by a broad range of neuromodulatory inputs, whereas expression of another immediate early gene, *Npas4*, is more tightly regulated by neuronal activity (Lin et al., 2008). We therefore tested whether GRP injections lead to similar changes in *Npas4* expression levels. Indeed, we found significantly increased *Npas4* expression following GRP injection in *Vip^+^* and glutamatergic cells, but not *Sst^+^* and *Pvalb^+^* cells (Fig. 4H and Fig. S4D_3_). Moreover, the cumulative distribution of normalized *Npas4* expression was significantly smaller when GRP was injected into *Vip-IRES-Cre;Grpr^fl/y^* mice compared to control mice (Kolmogorov-Smirnov: *Vip^+^* cells: p<0.0001 t=0.78; glutamatergic cells: p<0.0001 t=0.26, Fig. S4E_3_ and S4G), confirming GRP-mediated disinhibition of glutamatergic cells through VIP cell-specific GRPR signaling.

### ACx VIP cell activity encodes novel sounds and shocks during fear conditioning

The presence of *Grp^+^* cortically-projecting neurons in all areas of the thalamo-cortico-amygdala loop (Fig. 2E and 2K), a circuit central for the encoding of fear memories (Boatman and Kim, 2006), together with previous reports of stress- and fear-induced GRP release in the amygdala (Merali et al., 1998; Mountney et al., 2011) suggest that GRP mediates cortical modulation of sensory processing in the presence of aversive cues. An ideal behavioral paradigm to examine this possibility is auditory discriminatory fear conditioning that is based on foot shock-mediated changes in the encoding of auditory stimuli (Dalmay et al., 2019; Letzkus et al., 2011).

To first determine whether VIP cells in the primary ACx are recruited during this task we recorded bulk Ca^2+^- dependent fluorescence changes in GCaMP6s-expressing VIP cells using fiber photometry. To optimize comparison of VIP cell activity across mice and behavioral sessions, we tested the suitability of the AAV-CBA-DIO-GCaMP6s-P2A-mBeRFP construct for quantitative analysis of GCaMP fluorescence *in vivo*. We reasoned that if GCaMP and mBeRFP expression levels correlate and if mBeRFP fluorescence is entirely Ca^2+^-independent, mBeRFP red fluorescence, triggered by the same excitation wavelengths as that used for GCaMP, can be used as a reference to normalize GCaMP green fluorescence. In addition, the mBeRFP fluorescence can serve as a sensitive indicator of movement artifacts and patchcord detachment or entanglement. The right-shifted emission spectrum for mBeRFP in HEK 293T cells (473 nm excitation) compared to other frequently used red fluorophores (tdTomato and mRuby2 (Lam et al., 2012)), improved spectral separation from GCaMP fluorescence (Fig. S5A). Moreover, the addition of 2 mM Ca^2+^ and 10 µM ionomycin to HEK 293T cells expressing GCaMP strongly increased fluorescence when excited at 473 nm (t(5)=15.15, p=0.0001) as expected, but significantly decreased it when excited with 405 nm wavelength (a commonly used reference for photometric recordings that is incorrectly referred to as the isosbestic point for GCaMP; t(5)=-42.51, p<0.0001). In contrast, mBeRFP (t(5)=2.54, p=0.16), tdTomato (t(5)=0.94, p=0.78) and mRuby2 (t(5)=0.29, p=0.78) did not show any significant fluorescence changes (Fig. S5A). Importantly, the increase in GCaMP6s fluorescence upon Ca^2+^/ionomycin application was not different when comparing GCaMP6s alone to expression of CaMP6s-P2A-mBeRFP (t(10)=-0.12, p=0.91; n=6 wells each; Fig. 5A), indicating that mBeRFP is a Ca^2+^-independent fluorophore that does not quench or interfere with GCaMP6s fluorescence.

**Figure 5:**
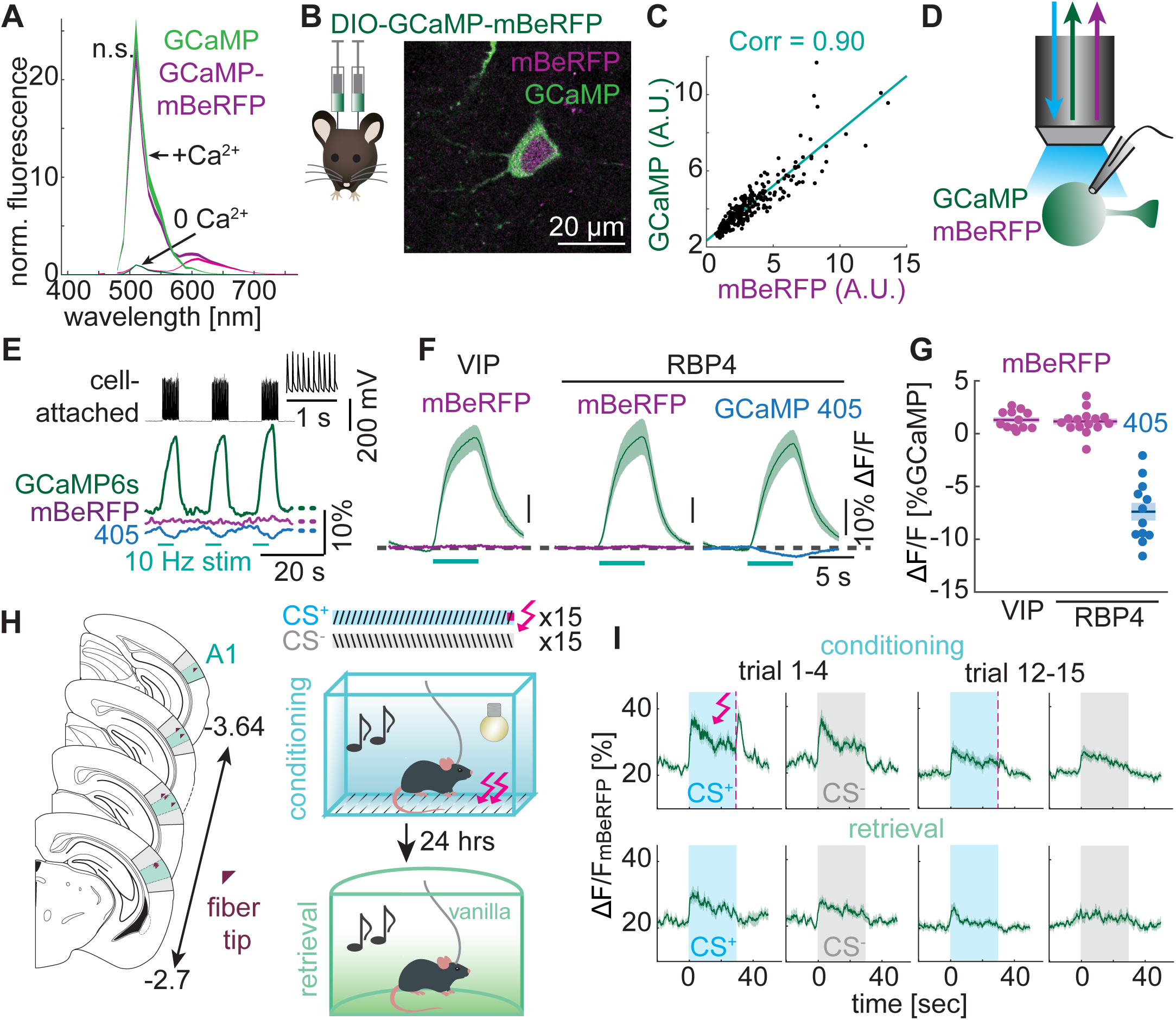
**ACx VIP cells encode novel sounds and shocks during fear conditioning** **A**, Fluorescence emission spectrum of GCaMP alone (green) and GCaMP-P2A-mBeRFP (magenta, pink) expressed in HEK 293T cells and measured with or without application of Ca^2+^/ionomycin. SEM (n=6 wells each). Excitation = 473 nm. Fluorescence was normalized to the maximum fluorescence in 0 Ca^2+^ in the range of 390 - 550 nm. Comparison of Ca^2+^-dependent fluorescence (390 – 550 nm): 2-sample t-test: t(10)=-0.12, p=0.91. **B**, Confocal image of a pyramidal cell after injection of AAV DIO-GCaMP-P2A-mBeRFP and AAV Cre into auditory cortex. **C**, Correlation of GCaMP and mBeRFP fluorescences in *Vip^+^* cells after injection of AAV DIO-GCaMP-P2A-mBeRFP into auditory cortex of *Vip-IRES-Cre* mice (linear regression and correlation coefficient in cyan). n=335 neurons from 3 injections sites. **D**, Schematic of experimental setup for analysis of action potential-dependent changes of GCaMP and mBeRFP fluorescence in acute brain slices during electrophysiological induction of spiking in VIP or L5 pyramidal (RBP4^+^) cells expressing AAV DIO-GCaMP-P2A-mBeRFP. **E**, Action potential bursts (each 5 s at 10 Hz) induced in an exemplary RBP4^+^ neuron through a cell-attached electrode in current clamp configuration (top) with GCaMP and mBeRFP fluorescence (473 nm excitation, *middle*) and GCaMP fluorescence (405 nm excitation, *bottom*). The inset shows individual spikes measured in the first second of the last burst. **F**, Average GCaMP and mBeRFP (473 nm excitation) and GCaMP (405 nm excitation) fluorescence changes (ΔF/F_0_). Data from cell-attached and whole-cell recordings were pooled. Dashed line: mean baseline fluorescence. Mean ± SEM from n≥3 mice each and n=12 VIP, 15 RBP4 GCAMP/mBeRFP, and 12 RBP4 GCaMP (405 nm) cells. **G**, Quantification of fluorescence changes shown in F normalized to GCaMP (473 nm) fluorescence change. Mean ± SEM. **H**, Schematics showing location of optical fiber tip for fiber photometric recordings of fluorescence of GCaMP-P2A-mBeRFP-expressing VIP cells in primary auditory cortex (*left*) obtained during fear conditioning and retrieval (*right*). Animals were exposed to up and down frequency sweeps (conditioning) one of which (CS^+^) was paired with electric shocks while the other sound (CS^-^) was presented unpaired. On the follow day (retrieval), both sounds were played in a new environment (shape, light and odor) and the effect on freezing levels assayed. **I**, GCaMP fluorescence changes (normalized to mBeRFP fluorescence) measured around presentation of conditioned (CS^+^, blue) and unconditioned sounds (CS^-^, grey) and shocks (dashed pink lines) early (trial 1-4) and late (trial 12-15) on the conditioning (top) or retrieval (bottom) day. Mean ± SEM across n=11 mice. See also Figure S5.

We tested AAV-CBA-DIO-GCaMP6s-P2A-mBeRFP expression and Ca^2+^ dependence in auditory cortical VIP and L5 pyramidal cells using Cre-dependent GCaMP6s-P2A-mBeRFP expression in *Vip-IRES-Cre* and Rbp4-Cre mice, respectively (Fig. 5B and S5B). Expression of GCaMP and of mBeRFP were highly correlated across cells (correlation coefficient: 0.90, n=335 cells in 3 *Vip-IRES-Cre* mice, Fig. 5C). Moreover, intrinsic and active physiological properties of VIP cells were not significantly different in GCaMP-P2A-mBeRFP- and GFP-expressing cells (Fig. S5C-E). To test the Ca^2+^ dependence of GCaMP and mBeRFP fluorescence in neurons, we activated GCaMP-mBeRFP-expressing neurons with 10 Hz electrical stimulation (5 sec duration, cell-attached or whole-cell mode, Fig. 5D-E). As expected, neuronal activity increased GCaMP fluorescence, whereas mBeRFP fluorescence was largely unaffected (mean mBeRFP ΔF/F_0_ relative to mean GCaMP ΔF/F_0_ during stimulation: VIP cells: 1.3 ± 0.3%; RBP4 cells: 1.1 ± 0.3% Fig. 5F-G). Of note, Ca^2+^ increases led to clear GCaMP fluorescence decreases when excited at 405 nm (-7.4 ± 0.8% relative to mean GCaMP ΔF/F_0_; Fig. 5F-G). These results establish mBeRFP as a non-toxic Ca^2+^-independent fluorophore that is useful for GCaMP fluorescence normalization.

To analyze VIP cell Ca^2+^ dynamics during fear conditioning, we acquired fiber-photometric recordings in GCaMP6s-P2A-mBeRFP-expressing VIP cells in the right primary auditory cortex (Fig. 5H and Fig. S5F). During the conditioning session, mice were fear conditioned to 15 complex sounds (conditioned stimulus, CS^+^, 30 frequency sweeps per sound) co-terminating with a brief foot shock, whereas 15 other complex sounds were presented without shocks (CS^-^, Fig. 5H). This fear conditioning paradigm, based on CS^+^ and CS^-^ frequency sweeps, is dependent on primary auditory cortex and regulated by cortical disinhibitory circuits (Dalmay et al., 2019; Letzkus et al., 2011). Fear retrieval 24 hrs after conditioning revealed that most mice had learnt the association between CS^+^ and foot shocks, as demonstrated by the increased freezing levels during CS^+^ presentation compared to baseline freezing (Fig. S5G-H). GCaMP fluorescence during CS^+^, CS^-^ and foot shocks was analyzed by subtracting autofluorescence (measured in 3 implanted mice without fluorophores) and subsequent normalization of GCaMP to mBeRFP fluorescence. We observed strong fluorescence increases in VIP cells in response to novel CS^+^, CS^-^ and aversive foot shocks (Fig. 5I). Responses to all three stimuli habituated strongly during conditioning and retrieval sessions (Fig. 5I). None of the responses were due to movement artifacts as demonstrated by stable mBeRFP fluorescence throughout sound and shock presentation (Fig. S5I). The strong initial activation upon aversive foot shocks, which triggered fast escape behavior, raise the possibility that VIP cell activity may be correlated with locomotion. However, neither cross-correlation of GCaMP fluorescence and speed (Fig. S5J) nor the trial-averaged fluorescence during movement initiation after bouts of freezing (Fig. S5K) during the initial phase of fear retrieval revealed strong GCaMP modulation by locomotion.

### GRPR signaling in the ACx enhances fear memories

To test whether GRP/GRPR signaling in VIP cells contributes to the modulation of fear memories, we used CRISPR/Cas9-mediated bilateral knock-out of *Grpr* in the ACx of male wild type mice. Out of four tested guide RNAs (sgRNAs) targeting *Grpr*, we selected the one (referred to as sgRNA-*Grpr1*) with the lowest predicted off-target cutting efficiency, high on-target cutting efficiency in cultured neuroblastoma cells (42%), and strong functional reduction of GRP-induced Ca^2+^ increases in ACx VIP cells when the CRISPR/Cas9 construct (AAV CMV-SaCas9-3xHA-U6-sgRNA (Tervo et al., 2016)) was co-expressed with GCaMP-P2A-mBeRFP (Fig. 6A-B). Thus, GRP-mediated Ca^2+^ increases in saCas9-expressing cells with sgRNA-*Grpr1* were significantly smaller compared to those with sgRNAs targeting the bacterial gene *lacZ* (ΔF/F_KCl_ : control (*lacZ*): 10.11 (12.1)%; *Grpr1*: -0.55 (4.3)%; U=25956; p<0.0001, n=602 and 298 cells in 10 and 9 slices for *lacZ* and *Grpr1* respectively; Fig. 6B; compare to the less efficient sgRNA referred to as *Grpr2*; Fig. S6A). Bilateral AAV CMV-SaCas9-3xHA-U6-sgRNA injections into the ACx of male wild type mice resulted in SaCas9-HA expression mainly in primary ACx with partial spread into dorsal and ventral ACx (Fig. 6C and S6B-C). Fear conditioning of control (*lacZ*) and KO (*Grpr1*) mice increased freezing during CS^+^ and CS^-^ throughout the conditioning session in both groups (2-way ANOVA, main effect of genotypes: CS^+^: p=0.90, F=0.01; CS^-^: p=0.78, F=0.08; no main interaction between genotype and stimulus number for CS^+^ and CS^-^, Fig. 6D). 24 hrs after conditioning, mice were subjected to auditory fear memory retrieval (Fig. 6E). Freezing levels during CS^+^, CS^-^ and after both sounds (30 sec period) were significantly reduced in KO mice compared to control mice (2-way ANOVA, main effect of genotype: p=0.0004, F=13.54, main effect of stimulus: p<0.0001, F=13.41; no significant interaction between genotype and stimulus; Fig. 6E-F and Fig. S6D), consistent with a function of GRPR signaling in enhancing auditory cortex-dependent memory formation.

**Figure 6:**
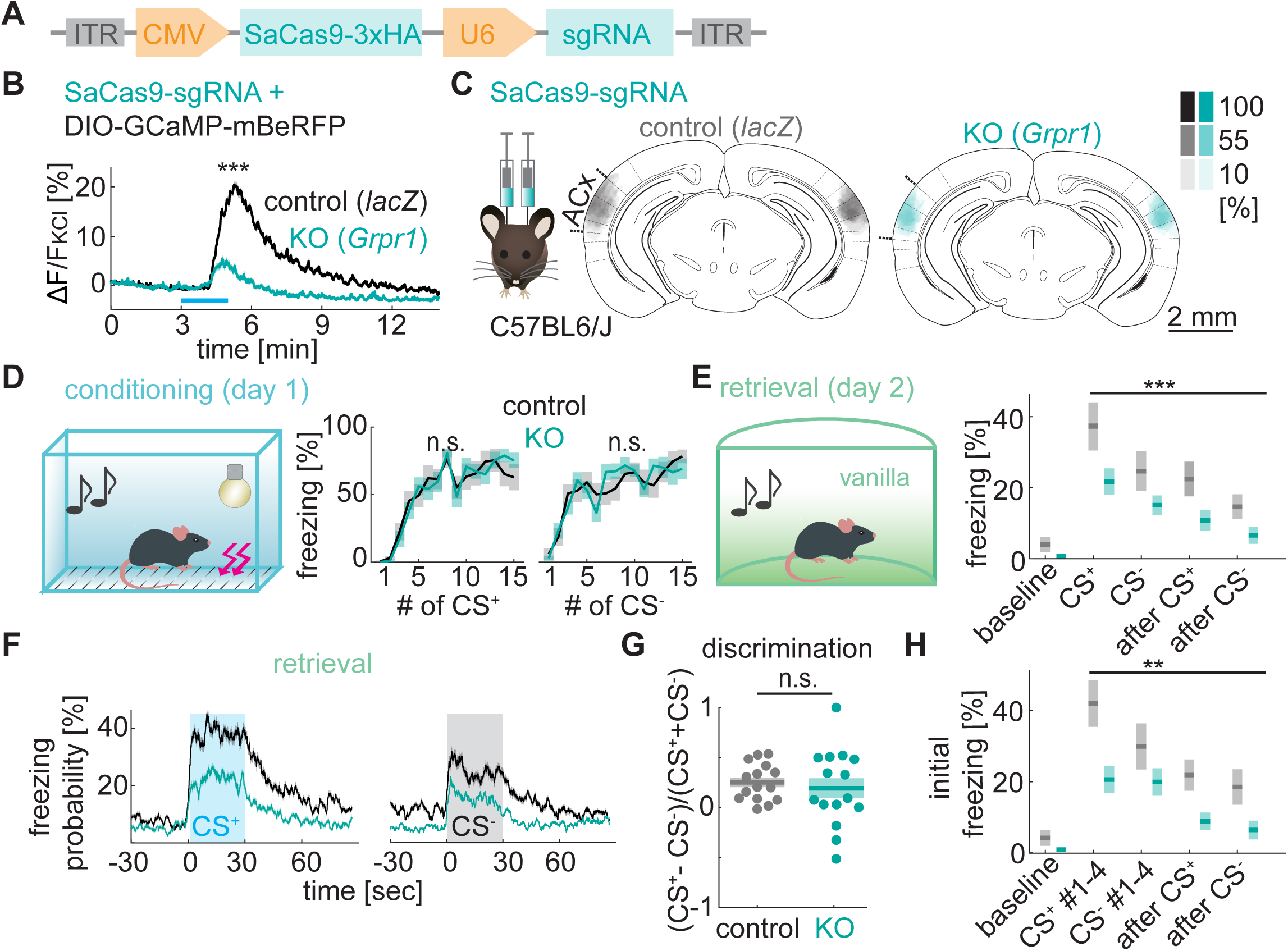
**GRPR signaling in the ACx enhances fear memories** **A**, Design of the plasmid for CRIPSR/Cas9-mediated KO of *Grpr* (here abbreviated as saCas9-sgRNA). **B**, GCaMP fluorescence changes measured in acute brain slices upon GRP application for VIP cells infected with AAV encoding saCas9-sgRNA targeting either *Grpr* (KO, *Grpr1*) or *lacZ* (control) (fluorescence is normalized to KCl fluorescence, mean ± SEM). Mann-Whitney U test: U=25956; p<0.0001, n=602 and 298 cells in 10 and 9 slices for *lacZ* and *Grpr1,* respectively. **C**: Quantification of bilateral saCas9-HA expression in mice with focal AAV injection into auditory cortex encoding sgRNA targeting *lacZ* (control, grey) or *Grpr* (KO, turquoise). See Figure S6 for analysis of expression in the whole brain. Color-code shows percentage of mice with saCas9-HA expression. **D**, Auditory fear acquisition, measured as the percentage of time spent freezing during presentation of 15 conditioned (CS^+^) and unconditioned (CS^-^) sounds on conditioning day in control and KO C57BL6/J mice (histology shown in C). Mean ± SEM. 2-way ANOVA, main effect of genotypes: CS^+^: p=0.90, F=0.01; CS^-^: p=0.78, F=0.08, no significant interaction of genotype and stimulus number. N=15 mice per group. **E**, Auditory fear memory retrieval, measured as the percentage of time spent freezing averaged across 15 presentations of CS^+^ and CS^-^ on the retrieval day. Mean ± SEM. 2-way ANOVA: main effect of genotype: p=0.0004, F=13.54, no significant interaction of genotype and stimulus. **F**, Time courses of average freezing probability across all CS^+^ and CS^-^ during fear memory retrieval for control (black) and KO (turquoise) mice. Mean ± SEM. **G**, Sound discrimination indices for control and KO mice measured during retrieval. Mean ± SEM. T-test for unequal variance: t(20.13)=0.53, p=0.60. **H**, Initial auditory fear memory retrieval, measured as the percentage of time spent freezing during first 4 CS^+^ and CS^-^ presentations during retrieval day. Mean ± SEM. 2-way ANOVA: main effect of genotype: p=0.005, F=8.67, no significant interaction of genotype and stimulus. See also Figure S6.

Neither the absolute freezing difference between CS^+^ and CS^-^ (2-sample t-test: t(28)=1.59, p=0.12) nor the discrimination index (difference between freezing level during CS^+^ and CS^-^ divided by the sum of both) were significantly different in control and KO mice (t-test for unequal variance: t(20.13)=0.53, p=0.60; Fig. 6G), suggesting that the behavioral effect is not due to an impairment in auditory discrimination. Since freezing levels did not change significantly over time (CS^+^: 2-way ANOVA, effect of genotype: p<0.0001, F=30.9, effect of stimulus number: p=0.85, F=0.61, no main interaction; CS^-^ effect of genotype: p=0.0001, F=15.71; effect of stimulus number: p=0.20, F=1.31, no main interaction; Fig. S6E), and freezing levels in KO mice were strongly reduced already during the first 4 CS^+^ and CS^-^ (2-way ANOVA: effect of genotype: p=0.005, F=8.67; effect of stimulus: p=0.237, F=1.43; no significant interaction), we conclude that the reduced freezing in KO mice is not due to accelerated fear extinction. Moreover, reduced freezing levels in KO mice were not purely a result of increased baseline freezing, since freezing levels were still significantly reduced in KO mice after subtraction of baseline freezing (2-way ANOVA: effect of genotype: p=0.029, F=5.05, effect of stimulus: p=0.024, F=5.4, no significant interaction; Fig. S6F).

Importantly, control and KO mice exhibited comparable locomotion during the conditioning session at baseline, during the first CS^+^ and CS^-^ and in response to the first foot shock (Fig. S6G-H: 2-way ANOVA, effect of genotype: p=0.93, F=0.01, no main interaction between genotype and stimulus), suggesting that the reduced freezing level in KO mice is not a result of different pain sensitivity or overall activity levels.

### Impaired fear memory in mice with conditional KO of GRPR in the ACx

To exclude that the observed behavioral effects were due to off-target cutting or other side effects of CRISPR/Cas9 expression, we repeated the behavioral experiments using conditional *Grpr* KO mice (Yu et al., 2017). To validate Cre-dependent knockout of *Grpr* in these mice we injected DIO-GCaMP-P2A- mBeRFP into the ACx of male *Vip-IRES-Cre* mice with VIP cell-specific knockout of *Grpr* (*Grpr^fl/y^;Vip-IRES-Cre*) or control mice (*Grpr^wt/y^;Vip-IRES-Cre*). GRP-induced Ca^2+^ increases in VIP cells of *Grpr^fl/y^;Vip-IRES-Cre* mice were strongly reduced compared to control mice (ΔF/F_KCl_: 16.9 ± 0.6% in ctrl vs. -1.0 ± 0.3% in KO mice, t(1178)=26.97, p<0.0001, n=655 cells in 14 slices and 525 cells in 11 slices respectively; Fig. 7A). To knockout *Grpr* specifically in the ACx for fear conditioning, we injected AAV hSyn-Cre-mCherry bilaterally into the ACx of *Grpr^fl/y^* (KO) or *Grpr^wt/y^* (ctrl) mice (Fig. 7B and S7A). Both groups of mice were exposed to fear conditioning and retrieval as above. Analysis of freezing during fear retrieval revealed significantly reduced freezing levels in *Grpr^fl/y^* (KO) mice upon and directly after presentation of CS^+^ and CS^-^ (2-way ANOVA: effect of genotype: p<0.0001, F=20.33, no significant interaction between genotype and stimulus; n=14 mice per group, Fig. 7C-D and S7B), with no significant effect on auditory discrimination (absolute freezing difference between CS^+^ and CS^-^: U=86, p=0.60). Importantly, the reduced freezing levels in KO mice were not caused by differences in genetic background of *Grpr^fl/y^* and *Grpr^wt/y^* mice, since freezing levels in uninjected *Grpr^fl/y^* mice were not different from their control littermates (2-way ANOVA: effect of genotype: p=0.84, F=0.04, no significant interaction between genotype and stimulus; n=16 mice per group, Fig. S7C-E), confirming that *Grpr* KO in the ACx reduces fear-induced freezing in a discriminatory auditory fear conditioning paradigm.

**Figure 7:**
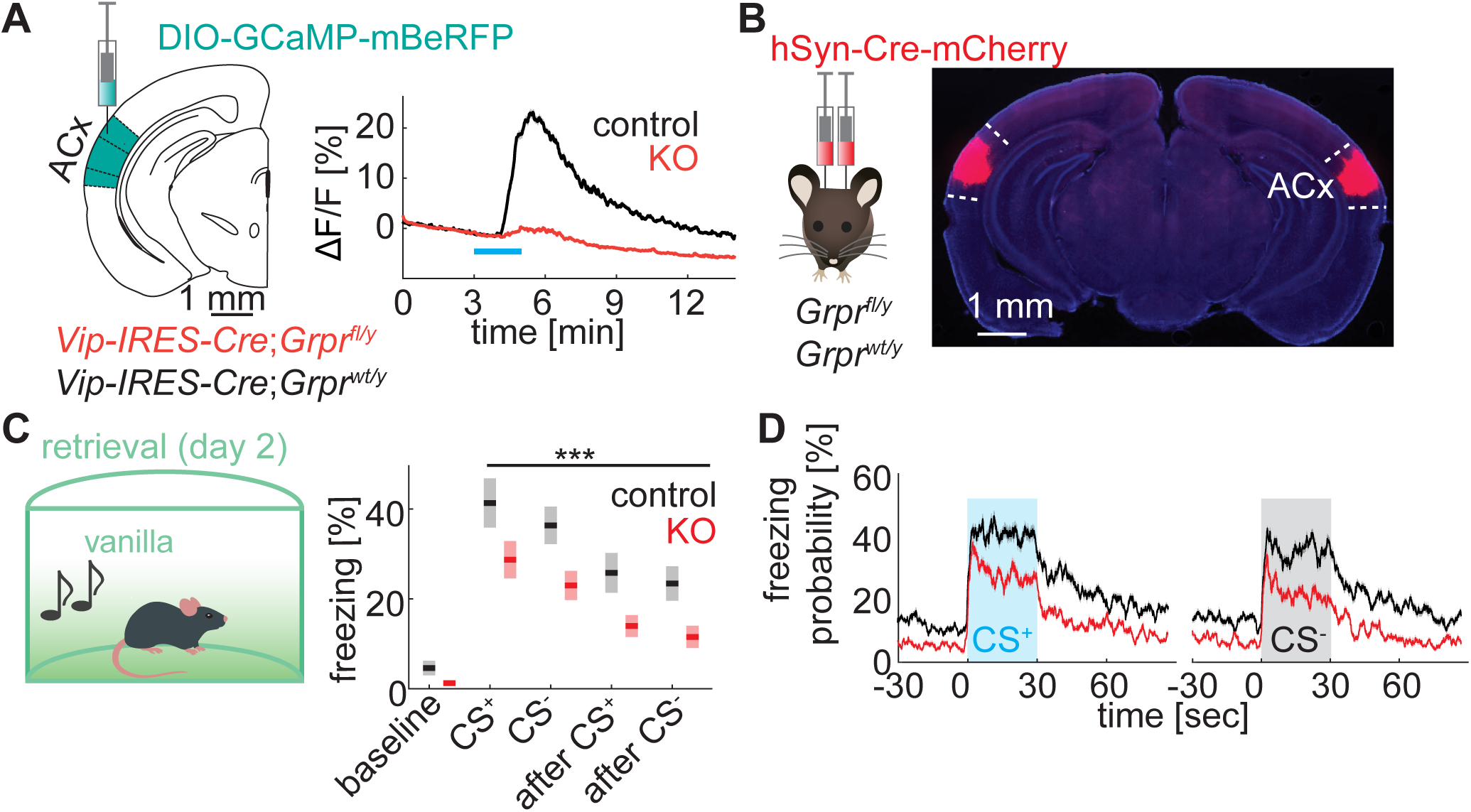
**Impaired fear memory in mice with conditional KO of GRPR in the auditory cortex** **A**, GCaMP fluorescence changes in VIP cells (normalized to KCl fluorescence) following GRP application in acute auditory cortical slices from control mice (*Vip-IRES-Cre;Grpr^wt/y^*) or mice lacking GRPR (*Vip-IRES-Cre;Grpr^fl/y^*). Mean ± SEM. Two-sample t-test: t(1178)=26.97, p<0.0001, n=655 cells in 14 slices (ctrl) and 525 cells in 11 slices (KO). **B**, Injection of AAV2 hSyn-Cre-mCherry into auditory cortex of *Grpr^wt/y^* or *Grpr^fl/y^* mice to locally knockout *Grpr*. Right: Epifluorescent image of exemplary injection sites. For quantification of expression levels across all mice see Figure S7. **C**, Time spent freezing during fear memory retrieval in *Grpr^wt/y^* (ctrl) and *Grpr^fl/y^* (KO) mice injected into auditory cortex with AAV encoding Cre-mCherry. Mean ± SEM. 2-way ANOVA: main effect of genotype: p<0.0001, F=20.33, no significant interaction of genotype and stimulus. N=14 mice per group. **D**, Time course of average freezing probability across all CS^+^ and CS^-^ presentations during fear memory retrieval. Mean ± SEM. See also Figure S7.

## Discussion

### Neuropeptidergic signaling for cortex-wide cell type-specific communication

Neuropeptidergic signaling has been suggested to interconnect cortical neurons and provide control over cortical homeostasis and plasticity (Smith et al., 2019). This hypothesis is supported by the cellular specificity of receptor-ligand expression patterns but has largely not been tested functionally. We here provide direct evidence that distinct peptidergic cell-to-cell communication channels, consistent with those predicted by mRNA expression patterns, exist and regulate cortex-dependent behaviors. We identify the bombesin-like peptide GRP as a neuropeptide that is expressed in distinct subsets of L2/3 and L6 cortical neurons and specifically targets VIP cells. Since in cerebral cortex GRP receptor expression is nearly entirely confined to VIP cells, GRP selectively depolarizes and activates these interneurons, and thereby disinhibits cortex, consistent with the synaptic architecture of the cortical microcircuitry (Karnani et al., 2016; Pfeffer et al., 2013; Pi et al., 2013). Although we limit our *in vivo* functional analysis to auditory cortex, we find that the specificity of GRP control over VIP cells and its ability to induce disinhibition are similar in all examined primary and higher-order cortical areas and thus provides a substrate for cortex-wide context-dependent modulation of signal transmission and plasticity. Furthermore, in addition to this intra-cortical communication channel, our data support a potential role for GRP in subcortical (thalamic and amygdalar) control of cortical circuits through the same VIP cell-specific targeting.

Unlike synaptic communication channels that have the ability to modulate individual cells through point-to-point communication, our data suggest that GRP acts on a larger scale through extrasynaptic diffusion. Thus, using a novel GRP sensor, we show that extrasynaptic GRP is viable for minutes, giving it the potential to signal through volume transmission and reach large populations of VIP cells. This mode of transmission for neuropeptides has been suggested several decades ago (Agnati et al., 1986) and is now thought to be a major communication mode for neuromodulators (Taber and Hurley, 2014).

Unfortunately, as our most sensitive grpLight sensors have apparent affinities (EC50) of ∼100-360 nM (Fig. S2.2F), they are unlikely to be able to detect endogenous GRP at physiologically-meaningful levels that occur at approximately 10% of this concentration.

### Cortical GRPR signaling enhances fear memories

Neuromodulators are thought to mediate context-dependent control over functional cortical circuits (Marlin et al., 2015; Nakajima et al., 2014; Polack et al., 2013; Smith et al., 2019). Most studies so far, however, focused on small-molecular neuromodulators such as acetylcholine and monoamines, which include dopamine, serotonin, and noradrenaline (Clark and Noudoost, 2014; Hasselmo, 2006; Polack et al., 2013; Tronel et al., 2004). We here provide evidence for a previously unknown peptidergic control mechanism of behaviorally-relevant cortical circuits. Thus, GRPR signaling normally enhances auditory fear memories as revealed by reduced freezing levels during fear retrieval when the highly selective GRP receptor (Battey and Wada, 1991; Kroog et al., 1995; Rozengurt, 1988) is deleted using conditional knockout and new CRISPR/Cas9-based tools. Our results are consistent with previous studies showing activation of and a contribution of primary auditory cortex to establishing auditory-associated fear memories (Dalmay et al., 2019; Letzkus et al., 2011). Our work extends these by providing the first evidence of VIP cell-specific and peptide-dependent modulation of fear memories in cortex.

The thalamo-amygdalo-cortical loop is central to auditory fear conditioning (Boatman and Kim, 2006). Interestingly, our data show that not only is GRP expressed in all regions in this loop but that it is found in the projection neurons that interconnect nodes of the circuit, forming GRP-expressing communication channels. Furthermore, previous findings that GRP is released in the amygdala upon acute stressors and by learnt aversive cues (Merali et al., 1998; Mountney et al., 2011) together with our own finding that GRP is expressed by amygdalo-cortical projections, suggests that GRP from amygdala projections is released in a context-dependent manner to disinhibit cortex as needed to strengthen memories in the presence of aversive or threatening cues. Moreover, previous studies found that trains of action potentials drive GRP release in the spinal cord (Pagani et al., 2019) and that amygdala neurons robustly increase activity upon foot shocks and fear-conditioned sounds (Grewe et al., 2017; Pelletier et al., 2005), thus supporting the hypothesis of context-dependent GRP signaling in auditory cortex.

Previous studies addressing GRP function in fear memory in the amygdala, the hippocampus and the prefrontal cortex yielded confounding results (Mountney et al., 2006, 2008; Roesler et al., 2003). This may be due, in part, to the use of pharmacological manipulations, such as injection of GRPR agonists, that are likely to desensitize receptors, as is evident in our and previous *in vitro* studies (Ally et al., 2003). For example, both agonist and antagonist infusions into infra-limbic cortex or basolateral amygdala decrease fear memories (Mountney et al., 2006, 2008). The use of genetic tools, including CRISPR/Cas9-mediated and conditional KO of GRPR as demonstrated here, will help disentangle the functions of GRPR in brain areas beyond cerebral cortex.

### VIP cells are recruited by novel, unexpected sounds and shocks

Although it is generally accepted that VIP cells are important regulators of cortical disinhibition and plasticity (Fu et al., 2014, 2015; Karnani et al., 2016; Pfeffer et al., 2013; Pi et al., 2013), evidence for memory and learning enhancement by activity of VIP cells is surprisingly scarce (Adler et al., 2019; Fu et al., 2015). Our results provide direct evidence for memory facilitation by VIP cell-specific signaling. Moreover, the cue-specific activation patterns of auditory cortex VIP cells during fear conditioning indicate encoding of novel sounds and shocks. VIP cell responses to sounds and aversive cues (air puffs) were observed previously in head-fixed animals (Pi et al., 2013), but it was unknown how VIP cells encode the changing valence of conditioned and unconditioned cues during fear conditioning. Interestingly, similar to amygdala VIP cells (Krabbe et al., 2019), we observed strong activation of VIP cells by novel unexpected shocks and habituation of VIP cell responses with learning of the sound-shock association. In contrast to amygdala VIP cells, however, we also observed strong responses to novel sounds, which later habituated, suggesting that the responses to unexpected sounds and shocks may both facilitate learning the sound-shock association. Interestingly, a distinct type of disinhibitory neurons in L1 of the auditory cortex also exhibit strong activation to shocks (information on habituation and sound responses are not available (Letzkus et al., 2011)), suggesting that both cells act synergistically to disinhibit pyramidal neurons (consistent with the decreased inhibition in pyramidal cells observed previously (Letzkus et al., 2011)), to facilitate shock-induced plasticity of sound responses.

### GRP-induced changes in signal transmission and plasticity

A remarkable feature of neuromodulator signaling is that each receptor can give rise to multiple intracellular signaling cascades with different temporal dynamics and functional implications (Belcheva and Coscia, 2002). We show here that GRP depolarizes VIP cells directly and selectively in several cortical areas, induces cell autonomous action potential-dependent and -independent cytosolic calcium transients, and increases expression of the immediate early gene *Fos*. We also observed GRP-induced calcium oscillations, which putatively depend on voltage-dependent calcium channels and Gαq-IP_3_-mediated calcium release from intracellular stores. Similar oscillatory and bombesin-induced calcium dynamics occur in insulin-producing hamster cells (Schöfl et al., 1996) and myenteric neurons (Simeone et al., 1995), and have been proposed to be important for differential regulation of gene expression and peptide secretion (Dolmetsch et al., 1998; Tse et al., 1993). To our knowledge, this is the first description of GRP-induced calcium oscillations in the brain.

Consistent with the circuit architecture that establishes VIP cells as the major disinhibitory cell type in multiple cortical areas (Karnani et al., 2016; Pfeffer et al., 2013; Pi et al., 2013), we found that GRP infusion inhibited SST and PVALB cells and disinhibited pyramidal cells. Whereas immediate disinhibition appears to require GRP levels sufficient to trigger action potentials in VIP cells, lower GRP concentrations may induce long-term changes in VIP cell excitability or GABA release through other G-protein dependent second messengers or beta-arrestin recruitment (Lefkowitz and Shenoy, 2005) with possible long-term effects on cortical disinhibition. Importantly, GRP infusion *in vivo* locally induced *Fos* and *Npas4* expression in VIP and pyramidal cells, which is one likely mechanism contributing to long-term plasticity and memory enhancement: Npas4 redistributes inhibitory inputs onto pyramidal cells (Bloodgood et al., 2013) and Fos facilitates long-term potentiation at hippocampal synapses (Fleischmann et al., 2003) and vesicle release (at least in *Drosophila* (Kim et al., 2009)).

### Towards a complete picture of neuromodulatory control of cortical circuits

The daunting task of unraveling the multitude of neuromodulatory effects on signal transmission and plasticity across all cortical cell types may be essential to understand how cortical function is modulated in different emotional/physiological states by the plethora of monoamine and neuropeptide receptors (Smith et al., 2019; Tasic et al., 2016, 2018). The heterogeneity of expression patterns and the diversity of receptors per cell type (Smith et al., 2019; Tasic et al., 2016, 2018) offer a glimpse into how intricate and versatile neuromodulator effects in the cortex can be, and highlight how much is left to be done to understand the effects of each neuromodulator on cortical circuits.

The analysis of transcriptomic data allows predictions of the effects of each neuromodulator receptor, especially for those that are confined to only one cell type, and will thus facilitate future studies. In fact, we show here that transcriptomic-based predictions are a powerful tool to motivate new studies of neuropeptides with unknown function. Our study shows modulatory control of a VIP cell-dependent disinhibitory microcircuit that facilitates auditory fear memories, consistent with our predicted disinhibitory effect of VIP cell-specific GRP receptors. Together with previous studies that revealed SST cell-specific effects of oxytocin in maternal and sociosexual behavior (Marlin et al., 2015; Nakajima et al., 2014), and SST cell-specific expression of a corticotropin-releasing hormone inactivating protein that induces anxiolytic effects (Li et al., 2016a), our study thus underlines the importance for future studies to investigate additional cell type-specific neuromodulatory communication channels and their impact on network activity and behavior.

Several local, intra-cortical neuropeptidergic interactions have been predicted (Smith et al., 2019). Importantly, our and previous studies on cortical neuropeptides (Li et al., 2016a; Marlin et al., 2015; Nakajima et al., 2014) highlight the importance of extending the analysis of cortical peptidergic communication channels to include subcortical areas. Thus, cortical oxytocin likely derives from hypothalamic subnuclei, whereas we show that GRP likely derives from amygdala and thalamic projection neurons additionally to local cortical neurons. The development of new peptide sensors based on the platform that we use here to develop the first GRP sensor, will facilitate future studies of region-specific peptide release from distinct local and long-range projecting cell classes. Our study points the way for future hypothesis-driven research motivated by cell type-specific expression of neuromodulatory receptors (Smith et al., 2019; Tasic et al., 2016, 2018), to unravel the functions of peptidergic and other modulatory communication channels, and their control over neuronal microcircuits and behaviors.

## ACKNOWLEDGEMENTS

We thank Byung Kook Lim for providing starting material for rabies virus production. We thank Zhou-Feng Chen for providing GRPR^fl/y^ mice. We thank Washington University for designing, testing and providing CRISPR/Cas9 constructs. We thank Zhihong Zhang and Liqun Luo for providing plasmids. We thank Ivo Spiegel and Mike Greenberg for sharing unpublished data for initial considerations of GPCRs. We thank Maree Webster (The Stanley Medical Research Institute) for providing fresh frozen tissue from human visual cortex. We thank the Neurobiology Department and the Neurobiology Imaging Facility for consultation and instrument availability that supported this work. This facility is supported in part by the Neural Imaging Center as part of an NINDS P30 Core Center grant #NS072030. We thank Dr. Barbara Caldarone and the Mouse Behavior Core for behavioral training and advice. This core is subsidized by the Harvard NeuroDiscovery Center. We thank members of the Sabatini lab for feedback on the manuscript. We thank Carolyn Marie Orduno Davis for assisting with grpLight *in vitro* screening and characterization.

The research was funded by grants from NIH (R35NS105107 (BS), U01NS115579 (LT and BS), NS072030 (P30 Core Center)), Q-FASTR, the Nancy Laurie Marks Foundation, Bertarelli, and by the German Research Foundation (DFG) postdoctoral fellowship, EMBO long-term fellowship, The Ellen R. and Melvin J. Gordon Center fellowship, the Brooks fellowship.

## AUTHOR CONTRIBUTIONS

S.M. and B.L.S. designed the experiments, discussed results and wrote the manuscript. S.M. coordinated the experiments and performed *in vitro* electrophysiology and imaging, histology, *in situ* hybridizations, analysis, immunohistochemistry, intracranial injections, behavior, photometry. E.N. performed behavior, intracranial injections, histology, *in situ* hybridizations, photometry, analyzed images and managed the mouse colony. G.O.M. and L.T. developed the GRP sensor. M.H. helped with cell culture imaging and designed some mBeRFP test experiments. A.C.P. performed cryostat sectioning, *in situ* hybridization and image analysis, maintained cell cultures and produced pseudotyped rabies virus. E.Q. performed intracranial injections, *in situ* hybridizations, cryostat sectioning, imaging and image analysis for CTB experiments. M.R.G. performed histology and image analysis. B.R. performed histology and intracranial injections. K.W.H. produced pseudotyped rabies virus. J.L. managed mouse colony.

## DECLARATION OF INTEREST

L.T and G.O.M are co-founders of Seven Biosciences.

## STAR METHODS

### Lead contact

Further information and requests for resources and reagents should be directed to and will be fulfilled by the Lead Contact, Bernardo L. Sabatini.

### Experimental model and subject detail

The following mouse lines were used in this study: Vip-EGFP (Tg(Vip-EGFP)JN37Gsat, MMRRC Cat# 031009-UCD, RRID:MMRRC_031009-UCD (Gong et al., 2003)), Sst-EGFP (FVB-Tg(GadGFP)45704Swn/J, The Jackson Laboratory, IMSR Cat# JAX:003718, RRID:IMSR_JAX:003718 (Oliva et al., 2000)), Pvalb-EGFP (CB6-Tg(Gad1-EGFP)G42Zjh/J, The Jackson Laboratory, IMSR Cat# JAX:007677, RRID:IMSR_JAX:007677 (Chattopadhyaya et al., 2004)), *Vip-IRES-Cre* (VIPtm1(cre)Zjh/J, The Jackson Laboratory, IMSR Cat# JAX:010908, RRID:IMSR_JAX:010908 (Taniguchi et al., 2011)), Rbp4-Cre mice (Tg(Rbp4-cre)KL100Gsat, MMRRC Cat# 031125-UCD, RRID:MMRRC_031125-UCD (Gong et al., 2003)), mice harboring floxed *Grpr* (B6;129S7-Grpr^tm1Zfc^/J, IMSR Cat# JAX:033148, RRID:IMSR_JAX:033148 (Yu et al., 2017)) and C57BL/6J mice (The Jackson Laboratory, IMSR Cat# JAX:000664, RRID:IMSR_JAX:000664). Transgene expression in VIP cells was previously characterized in the hippocampus (Tyan et al., 2014).

All mice except for Sst-EGFP mice were backcrossed to C57BL/6J mice for at least 6 generations. Mice of either sex were used at postnatal days 18 – 100.

Human tissue for FISH was obtained from The Stanley Medical Research Institute.

Animals used for *in vitro* experiments were group-housed, animals used for behavioral experiments were single-housed 4 to 7 days before start of the behavior. All mice were kept on a 12 h light/dark cycle. All experiments were conducted during the light phase of the schedule.

All procedures were performed in accordance with protocols approved by the Harvard Standing Committee on Animal Care following guidelines described in the U.S. National Institutes of Health Guide for the Care and Use of Laboratory Animals.

### Method details

#### Plasmids

For the generation of the pAAV-CBA-DIO-GCaMP6s-P2A-mBeRFP-WPRE-pA plasmid, mRuby2-P2A was cut out from pAAV-CBA-FLEX-mRuby2-P2A-GCaMP6s via AgeI and NheI restriction sites and replaced by de novo synthetized Kozak-sequence via AgeI and NheI cloning sites: GCTAGCCATACCATGATGATGATGATGATGAGAACCCATGGTGGCACCGGT. The STOP codon of GCaMP6s was then replaced by de novo synthetized GSG-P2A-mBeRFP via PCR cloning (Clone EZ): GGAATTCTTATTAAAGTTTGTGCCCCAGTTTGCTAGGGAGGTCGCAGTATCTGGCCACAGCCACCTCGTGCTGCTCGACGGAGGTCTCTTTGTCGGCCTCCTTGATTCTTTCCAGTCTTCTGTCCACATAGTAGACGCCGGGCATCTTGAGGTTCTTAGCGGGTTTCTTGGATCTGTATGTGGTCTTGGCGTTGCAGATCAGGTGGCCCCCGCCCACGAGCTTCAGGGCCATGTAGTCTCTGCCTTCCAGGCCGCCGTCAGCGGGGTACAGCATCTCGGTGCTGGCCTCCCAGCCGAGTGTTTTCTTCTGCATCACAGGGCCGTTGGATGGGAAGTTCACCCCTCTGATCTTGACGTTGTAGATGAGGCAGCCGTCCTGGAGGCTGGTGTCCTGGGTAGCGGTCAGCACGCCCCCGTCTTCGTATGTGGTGGATCTCTCCCATGTGAAGCCCTCAGGGAAGGAC TGCTTAAAGAAGTCGGGGATGCCCTGGGTGTGGTTGATGAAGGTCTTGCTGCCGTACATGAAGCTGGTAGCCAGGATGTCGAAGGCGAAGGGGAGAGGGCCGCCCTCGACCACCTTGATTCTCATGGTCTGGGTGCCCTCGTAGGGCTTGCCTTCGCCCTCGGATGTGCACTTGAAGTGGTGGTTGTTCACGGTGCCCTCCATGTACAGCTTCATGTGCATGTTCTCCTTAATCAGCTCTTCGCCCTTAGACACCATAGGACCGGGGTTTTCTTCCACGTCTCCTGCTTGCTTTAACAGAGAGAAGTTCGTGGCTCCGGATCC.

The sequence of the final construct was verified by sequencing.

pRSET-BeRFP plasmid was given to us by Zhihong Zhang (Yang et al., 2013). pAAV-CBA-FLEX-mRuby2-GSG-P2A-GCaMP6s-WPRE-pA was a gift from Tobias Bonhoeffer and Mark Huebener and Tobias Rose (Addgene plasmid # 68717) (Rose et al., 2016).

pcDNA3.1-mBeRFP was generated by cloning de novo synthetized mBeRFP into the pcDNA3.1(+) vector via BamHI-EcoRI cloning sites: GCCACCATGGTGTCTAAGGGCGAAGAGCTGATTAAGGAGAACATGCACATGAAGCTGTACATGGAGGGCACCGTGAACAACCACCACTTCAAGTGCACATCCGAGGGCGAAGGCAAGCCCTACGAGGGCACCCAGACCATGAGAATCAAGGTGGTCGAGGGCGGCCCTCTCCCCTTCGCCTTCGACATCCTGGCTACCAGCTTCATGTACGGCAGCAAGACCTTCATCAACCACACCCAGGGCATCCCCGACTTCTTTAAGCAGTCCTTCCCTGAGGGCTTCACATGGGAGAGATCCACCACATACGAAGACGGGGGCGTGCTGACCGCTACCCAGGACACCAGCCTCCAGGACGGCTGCCTCATCTACAACGTCAAGATCAGAGGGGTGAACTTCCCATCCAACGGCCCTGTGATGCAGAAGAAAACACTCGGCTGGGAGGCCAGCACCGAGATGCTGTACCCCGCTGACGGCGGCCTGGAAGGCAGAGACTACATGGCCCTGAAGCTCGTGGGCGGGGGCCACCTGATCTGCAACGCCAAGACCACATACAGATCCAAGAAACCCGCTAAGAACCTCAAGATGCCCGGCGTCTACTATGTGGACAGAAGACTGGAAAGAATCAAGGAGGCCGACAAAGAGACCTCCGTCGAGCAGCACGAGGTGGCTGTGGCCAGATACTGCGACCTCCCTAGCAAACTGGGGCACAAACTTTAAT

CRISPR/Cas9 plasmids were cloned at the Genome Engineering and iPSC Center (GEiC) at Washington University. Single guide RNAs (sgRNAs) were designed in Benchling, and in collaboration with the GEiC. The four sgRNAs that were tested *in vitro* at the GEiC targeted the second exon (first exon within coding sequence, within first 285 nucleotides following the start codon) of *Grpr* (NM_008177.3) and were predicted to have high on- and low off-target cutting in Benchling and at the GEiC.

sgRNA-*Grpr1*: GATGATAAGCCCATAAACTGCNNGRRT

sgRNA-*Grpr2*: AACGACACCTTCAATCAAAGTNNGRRT

sgRNA-*Grpr3*: CAGGGATGACATAGATGAAGCNNGRRT

sgRNA-*Grpr4*: CTGCTGGTGACATGCGCCCCTNNGRRT

The control sgRNA was desgined in Benchling, targeted at the bacterial *lacZ* gene. We selected a sequence with no predicted on- and off-target cutting for *Mus musculus* coding DNA using Benchling analysis. sgRNA-*lacZ*: CATCGCGTGGGCGTATTCGCA

sgRNAs were cloned into pAAV-CMV-Kozak-NLS-saCas9-NLS-3xHA-Tag-pA-U6-sgRNA via BsaI cloning sites. This plasmid was a gift from Feng Zhang (Addgene plasmid # 61591) (Ran et al., 2015).

sgRNA-*Grpr1* and sgRNA-*Grpr2* were tested in vivo because of their high on-target cutting frequencies in cultured mouse neuroblastoma (N2a) cells (42% and 24% respectively) as confirmed by next-generation sequencing, and because of their low predicted off-target cutting. sgRNA-*Grpr2* excluded because of too low functional *Grpr* KO efficiency based on GCaMP6s imaging. sgRNA-*Grpr3* had high on-target cutting (64%), but was excluded because of high predicted off-target cutting. sgRNA-*Grpr4* was excluded because of too low on-target cutting efficiency in N2a cells (4%).

pcDNA3-mRuby2 was a gift from Michael Lin (Addgene plasmid # 40260) (Lam et al., 2012). pAAV-EF1a-FAS-TdTomato (Addgene plasmid # 37092) (Saunders et al., 2012).

pAAV-CAG-GCAMP6s-WPRE-SV40 was a gift from Douglas Kim and GENIE Project (Addgene plasmid # 100844) (Chen et al., 2013).

pAAV-EF1a-Cre was a gift from Karl Deisseroth (Addgene plasmid # 55636) (Fenno et al., 2014).

pAAV CAG-DO-TC66T-mCherry plasmid was generated by restriction enzyme digest of pAAV-CAG-Flex-TC66T with SalI and AscI to cut out the inverted reading frame encoding TC66T-mCherry. The endings of TC66T-mCherry were then mutated to reverse the AscI and SalI restriction enzyme sites and the new insert SalI-TC66T-mCherry-AscI ligated into the original vector as a non-inverted reading frame. The CAG-Flex-TC66T plasmid was a gift from Liqun Luo (Addgene plasmid # 48331) (Miyamichi et al., 2013).

pAAV-CAG-FLEX-oG-WPRE-SV40pA was a gift from Edward Callaway (Addgene plasmid # 74292) (Kim et al., 2016).

pAAV-CBA-DIO-GCaMP6s-P2A-mBeRFP-WPRE-pA, pAAV CAG-DO-TC66T-mCherry, pRSET-BeRFP and the CRISPR/Cas9 constructs will be deposited in the online repository Addgene (addgene.org).

#### Immunohistochemistry

Mice were transcardially perfused with 4% paraformaldehyde (PFA). Coronal sections were cut at 50 μm thickness on a Leica VT 1000S Vibratome and washed with phosphate buffered saline (PBS). Free-floating sections were permeabilized and blocked for 1 hr with PBS containing 5% NGS and 0.2% Triton X-100. Incubation of the sections with primary antibodies was performed for 24 hrs at 4°C in fresh PBS containing 5% NGS and 0.2% Triton X-100. For double-labeling experiments both primary antibodies were incubated simultaneously. Sections were washed with PBST 5x and incubated for 1 hr with 1:500 Alexa 647-goat anti-rabbit IgG (Invitrogen). After repeated washing with PBST and PBS, the sections were mounted on glass slides. Pictures were taken using a virtual slide microscope (Olympus VS120), a laser scanning confocal microscope (Leica TCS SP8) or a fluorescence microscope with structured illumination (Keyence, BZ-X710).

#### Primary Antibodies

1:5000 rabbit-anti Fos (Synaptic Systems Cat# 226 003, RRID:AB_2231974) 1:1500 rabbit anti HA-Tag (C29F4, Cell Signaling Technology Cat# 3724, RRID:AB_1549585) 1:500 chicken anti-GFP (Abcam Cat# ab13970, RRID:AB_300798)

#### Fluorescent *in situ* hybridization

Mice were deeply anesthetized with isoflurane and decapitated, and their brains were quickly removed and frozen in Tissue Tek OCT compound (VWR, Radnor PA) on dry ice. Brains were cut on a cryostat (Leica CM 1950) into 20 μm thick coronal sections, adhered to SuperFrost Plus slides (VWR, Radnor PA), and immediately refrozen. Samples were fixed in 4% paraformaldehyde for 15 min at 4 degrees, processed according to RNAscope Fluorescent Multiplex Assay manual (Advanced Cell Diagnostics, Newark CA), and coverslipped with ProLong antifade reagent (Molecular Probes, Eugene, OR). *Gad1* and *Gad2* probes were combined in one channel. *Slc17a6,7,8* were combined in one channel. *Sst* probe and *RabV-gp1* probe were diluted 1:1 and 1:200 respectively to avoid bleed through into other fluorescnece channels. For protease treatment, slices were incubated in Protease III for 20 min. CTB-labeled slices were treated with protease III for only 10 min to preserve endogenous CTB fluorescence. The following probes were used: Mm-*Fos* (#316921-C3), Mm-*Sst* (#404631-C1 and C3), Mm-*Pvalb* (#421931-C2 and C3), Mm-*Vip* (#415961-C1 and C3), Mm-*Gad1* (#400951-C3), Mm-*Gad2* (#439371-C3), Mm-*Slc17a6* (#319171-C2), Mm-*Slc17a7* (#416631-C2), Mm-*Slc17a8* (#431261-C2), Mm-*Grp* (#317861-C1 and C3), Mm-*Grpr* (#317871-C2), V-*RabV-gp1* (#456781-C2 and C3 targeting rabies virus nucleoprotein N), *Cre* (#312281), Mm-*Crh* (#316091), Mm-*Npas4* (#423431), *oG* (#519441-C2).

For FISH on human visual cortex slices, 14 μm thick fresh frozen sections were used. For protease treatment, slices were incubated in Protease IV for 40 min. Tissue was obtained from a white male, 53 years old, who died of a heart attack without known diseases. The following probes were used: Hs-*GRPR* (#460411-C1 and C3), Hs-*GRPR*-O1 (#465271-C1), Hs-*VIP* (#452751-C2), Hs-*GAD1* (#404031-C3).

Images were taken at a Leica SP8 X confocal microscope using a 63x 1.4 NA oil immersion objective (Harvard NeuroDiscovery Center), at a pixel size of 180 nm (or 240 nm for rabies tracing and *Fos* and *Npas4* analysis) and an optical section of 0.9 μm.

For cell type-specific quantification of *Grp* and *Grpr* expression levels, ROIs were drawn semi-automatically and manually optimized in ImageJ (RRID:SCR_003070). *Grp* and *Grpr* images were thresholded using either RenyiEntropy or a manually set threshold, and the percent coverage of each ROI with *Grp* and *Grpr* puncta was quantified in ImageJ. Follow-up analysis was done in MATLAB (RRID:SCR_001622).

#### Stereotactic AAV injections

5-8 weeks old mice were surgerized. Anesthesia was induced with 5% isoflurane and maintained with 1-2.5% isoflurane. Ketoprofen (5-10 mg/kg) was given subcutaneously prior to incision. For injections, a small craniotomy (∼ 1 mm diameter) was made using the following coordinates (distance from bregma [mm] / distance from midline [mm] / depth [mm] / angle [°]):

Motor cortex: 0.6 / 1.5 / 0.7

ACx: -3.25 / as lateral as possible / 0.9

A glass micropipette was inserted through a small durotomy for virus delivery. The pipette was held in place for 10 min. 400 nl AAV were injected at 40 nl/min using a UMP3 microsyringe pump (World Precision Instruments, Sarasota FL). The pipette was held in place for another 10 min after the end of the injection. The pipette was then slowly retracted. The scalp incision was sutured, and post-surgery analgesics were given to aid recovery for three days (5-10mg/kg ketoprofen injected subcutaneously every 24 hours). AAVs were allowed to express for at least 3 weeks before the mice were used for experiments.

The following AAVs and titers were used:

AAV2/2 pX601-AAV-CMV::NLS-saCas9-NLS-3xHA-bGHpa;U6::BsaI-sgRNA-*Grpr1* (from Boston Children’s hospital viral core, BCH): 1.01*10^13 GC/ml

AAV2/2 pX601-AAV-CMV::NLS-saCas9-NLS-3xHA-bGHpa;U6::BsaI-sgRNA-*Grpr2* (from Boston Children’s hospital viral core, BCH): 1.22*10^13 GC/ml

AAV2/2 pX601-AAV-CMV::NLS-saCas9-NLS-3xHA-bGHpa;U6::BsaI-sgRNA-*lacZ* (from BCH): 1.67*10^12

AAV2/2 hSyn-Cre-mCherry (from UNC Vector Core): 1.55*10^12 GC/ml AAV2/DJ CBA-DIO-GCaMP6s-P2A-mBeRFP (from BCH): 1*10^12 GC/ml AAV2/9 CAG-DIO-oG (BCH): 1*10^11 GC/ml

AAV2/9 CAG-DIO-TC66T-mCherry (BCH): 1*10^11 GC/ml AAV2/8 CAG-DO-TC66T-mCherry (BCH): 1*10^11 GC/ml

AAV2/9 CAG-Flex-EGFP-WPRE-bGH (Penn Vector Core): 9.3*10^12 GC/ml AAV2/1 hSyn grpLight1.2 (BCH): 10^9 GC/ml (neuronal culture)

AAV2/1 hSyn grpLight1.3 (BCH): 10^9 (neuronal culture); 1.6*10^13 GC/ml (*in vivo*) AAV2/1 hSyn grpLight1.3ER (Vigene Biosciences): 10^9 GC/ml (neuronal culture)

AAVs from BCH were purified by iodixanol gradient purification and ultracentrifugation at 48krpm for 1 hr. To avoid leak expression of Cre-dependent AAVs, all Cre-dependent AAVs were first injected into wild type mice at various titers to determine the titer at which no expression was detected in the control mice in the absence of Cre-recombinase.

#### Retrograde tracer injection (CTB)

6 to 8 weeks old wild type mice were injected into the ACx, motor cortex or auditory thalamus with 25-100 nl CTB 647 or CTB 555 (4 μg/μl, Molecular Probes, Eugene, OR). The surgery was as described above with the following changes: CTB was injected with a flow rate of 20 nl/min. The coordinates for the auditory thalamus were (in mm) -3.1 posterior to bregma, 2.2 lateral to midline, 3.3 deep. To avoid CTB leak into cortex after injections into the auditory thalamus, the pipette was retracted from the brain at around 2.75 μm per second.

10 days after injection, mice were sacrificed, and the brains used for *in situ* hybridization as described above. Images were taken at a Leica SP8 X confocal microscope using a 63x 1.4 NA oil immersion objective (Harvard NeuroDiscovery Center), a pixel size of 180 nm and an optical section of 0.9 μm.

#### Rabies virus tracing

EnvA-pseudotyped, glycoprotein-deleted rabies virus carrying nuclear localized EGFP transgene (SADΔG-H2B:EGFP(EnvA)) was generated in house, using starting materials from Byung Kook Lim (UCSD). The recombinant rabies viruses were generated as described previously (Mandelbaum et al., 2019) using protocols similar to those established previously (Wickersham et al., 2010). In short, HEK 293T cells (ATCC Cat# CRL-11268, RRID:CVCL_1926) were transfected with pSPBN-SADΔG-H2B:EGFP (Mandelbaum et al., 2019), pTIT-B19N, pTIT-B19G, pTIT-B19L and pCAGGS-T7. Virions were then retrieved from the supernatant, amplified in BHK-B19G cells and concentrated through a serious of filtration and centrifugation steps. Pseudotyping was performed by infecting BHK-EnvA cells with virions from the previous step. Pseudotyped rabies virus titer was estimated based on serial dilution method (Osakada and Callaway, 2013), counting infected H2B:EGFP^+^ HEK 293T-TVA800 cells and quantified as infectious units per ml (IU/ml). Pseudotyped rabies virus was used at a titer of approximately 1 x 10^9 IU/ml. For quality control, HEK 293T cells were infected with pseudotyped rabies virus at serial dilutions and H2B:EGFP^+^ cells counted. The rabies virus batch used in this study had a leak of less than 2 x 10^3^ IU/ml. Aliquots were stored at -80°C.

For transsynaptic retrograde tracing, 6-8 weeks old *Vip-IRES-Cre* mice were injected with 200 nl AAV2/9 CAG-DIO-TC^66T^-mCherry and AAV2/9 CAG-DIO-oG (10^11 GC/ml each) into the auditory or motor cortex. The surgery was as described above. 3 weeks later, 200 nl SADΔG-H2B:EGFP(EnvA) was injected into the ACx or motor cortex as described above. Mice were sacrificed 7 days later and brains were sectioned on a cryostat for subsequent *in situ* hybridization or perfused and sliced on a vibratome for subsequent epifluorescent and confocal imaging.

The following control injections were performed to verify specificity of rabies tracing:

1. 200 nl SADΔG-H2B:EGFP(EnvA) was injected into the cortex to confirm specific specificity for TC^66T^- expressing cells. No labeling was detected.
2. 200 nl AAV2/8 CAG-DO-TC^66T^-mCherry (Cre-off version of TC^66T^, see Plasmids) and AAV2/9 CAG-DIO-oG (10^11 GC/ml each) were injected into the cortex of wild type mice followed by SADΔG-H2B:EGFP(EnvA) injections to confirm that oG expression was Cre-dependent in the presence of strong TC^66T^ and SADΔG-H2B:EGFP(EnvA) expression.
3. 200 nl AAV2/9 CAG-DIO-TC^66T^-mCherry and AAV2/9 CAG-DIO-oG (10^11 GC/ml each) were injected into the cortex of wild type mice followed by SADΔG-H2B:EGFP(EnvA) injections to confirm that TC^66T^ expression and SADΔG-H2B:EGFP(EnvA) transfection were Cre-dependent.

For *in situ* hybridization, every 4^th^ section (20 μm thick) was used to identify and quantify starter cells defined by the expression of *Cre*, *oG* and the rabies-specific nucleoprotein N (*in situ* probe V-*RABV-gp1*). Endogenous mCherry expression (from AAV2/9 CAG-DIO-TC^66T^-mCherry) was faint and diffuse due to tissue processing and could be clearly distinguished from in situ labeling (mCherry and *Cre* were imaged in the same channel). Endogenous EGFP fluorescence from SADΔG-H2B:EGFP(EnvA) rabies virus was strongly reduced due to tissue processing. The *in situ* probe V-*RABV-gp1* was therefore used to optimize detection of rabies virus. V-*RABV-gp1* and EGFP were imaged in the same channel. Every 4^th^ section was used to identify and quantify retrogradely labeled cells and *Grp* coexpression in the auditory thalamus using the *in situ* probes V-*RABV-gp1* and Mm-*Grp.* Every 8^th^ cryostat section was used to identify and quantify retrogradely labeled cells and *Grp* coexpression in the cortex surrounding the injection site. Due to the high number of retrogradely labeled cells surrounding the injection site, we found this sampling rate to be adequate to analyze a sufficiently large number of cells. We normalized the number of retrogradely labelled cells to the number of detected starter cells.

The V-*RABV-gp1* probe was diluted 1:100 or 1:200 in probe diluent to avoid bleed through into other channels. The whole auditory and motor cortices were screened for starter cells. *Grp* colabeling in retrogradely labeled cells was quantified in sections from 800 μm anterior to 800 μm posterior to the injection site, and in the entire auditory thalamus.

Images were taken at a Leica SP8 X confocal microscope using a 63x 1.4 NA oil immersion objective (Harvard NeuroDiscovery Center), at a pixel size of 240 nm and an optical section of 0.9 μm.

Cells were counted manually.

#### Immediate early gene expression analysis

200 nl full-length mouse GRP (Phoenix Pharmaceuticals #027-40, 3 µM in NRR) or NRR (in mM: 135 NaCl, 5.4 KCl, 5 HEPES, 1.8 CaCl_2_, pH 7.2 adjusted with KOH, sterile-filtered with 0.2 μm pore size) were injected into the right motor cortex of 6-8 week old male C57BL/6J, *Vip-IRES-Cre;Grpr^wt/y^* or *Vip-IRES-Cre;Grpr^fl/y^* mice. The surgery was performed as described above for AAV injections. Mice woke up from anesthesia within a few minutes after the end of the surgery and were allowed to move freely in their recovery chamber. 45 min after GRP or NRR were injection, mice were re-anesthetized, brains quickly dissected and frozen in OCT for *in situ* hybridization as described above. For immunostaining, mice were perfused 1 1/2 hours after, and 50 μm thick coronal slices were cut on a vibratome for anti-FOS immunostaining (see protocol above).

Images of *in situ* hybridized tissue were taken at a Leica SP8 X confocal microscope equipped with a 63x 1.4 NA oil immersion objective (Harvard NeuroDiscovery Center), at a pixel size of 240 nm and an optical section of 0.9 μm. Confocal images of 460-470 μm width spanning all cortical layers were taken at a fixed distance from the midline from both hemispheres (2-3 slices per mouse, surrounding the injection site). ROIs were drawn manually around all stained *Vip^+^*, *Sst^+^*, *Pvalb^+^* and *Slc17a6/Slc17a7^+^* cells in ImageJ. *Fos* and *Npas4* images were thresholded manually. Slices were discarded if left and right hemisphere exhibited different background fluorescence. Percent coverage of each ROI with *Fos* or *Npas4* puncta was quantified in ImageJ.

*Fos* and *Npas4* expression levels in the left (uninjected) hemisphere were used as a baseline to account for variability in staining intensity and baseline *Fos* and *Npas4* expression. Data from both hemispheres are shown in most figures. Expression levels were defined as cell area covered with FISH labeling. Mean expression across cortical depth was calculated as average expression across sliding windows of 150 µm width for each slice. Cumulative distributions of *Fos* and *Npas4* expression in the right hemisphere were calculated by normalizing the expression of all cells of a defined cell type in the right hemisphere to the mean expression of *Fos* or *Npas4* in the same cell type in the left hemisphere (for each slice).

Due to the high excitability (and *Fos* induction) of ACx circuits upon damage of microvessels following glass pipette insertion, we limited our analysis to the motor cortex.

#### RNA sequencing analysis

Visual cortex RNA sequencing data were downloaded from the Allen Brain Institute: https://portal.brain-map.org/atlases-and-data/rnaseq (Tasic et al., 2018) http://casestudies.brain-map.org/celltax#section_explorea (Tasic et al., 2016) Gene counts were normalized to counts per million (CPM). The two datasets were then screened for genes whose expression was correlated to *Vip* express with a correlation coefficient of >0.5.

#### Electrophysiological recordings

For *in vitro* patch-clamp recordings, mice were deeply anesthetized with inhaled isoflurane, and transcardially perfused with ∼30 ml ice-cold sucrose solution oxygenated with carbogen gas (95% O_2_, 5% CO_2_, pH 7.4). Mice were decapitated and brains removed. 300 μm thick sections were cut on a Leica VT 1000S vibratome in ice-cold oxygenated sucrose solution containing (in mM) 252 sucrose, 3 KCl, 1.25 Na_2_H_2_PO_4_, 24 NaHCO_3_, 2 MgSO_4_, 2 CaCl_2_, 10 glucose. Coronal slices were used for all experiments. Slices were incubated in oxygenated Ringer’s extracellular solution containing (in mM) 125 mM NaCl, 25 mM NaHCO_3_, 1.25 mM NaH_2_PO_4_, 2.5 mM KCl, 2 mM CaCl_2_, 1 mM MgCl_2_, 25 mM glucose at 32°C for ∼15 min, and subsequently at RT until used for recordings. Whole-cell patch-clamp recordings were performed at 30-32°C using pipettes pulled from borosilicate glass capillaries with resistances of 3-4 MΩ. Sections were continuously perfused with oxygenated extracellular solution. Cells were visualized by an upright microscope equipped with Dodt gradient contrast and standard epifluorescence.

All electrophysiological recordings were acquired using Multiclamp 700B amplifier (Molecular Devices) MultiClamp Commander and MTTeleClient for telegraphs. MATLAB (RRID:SCR_001622) was used to control current/voltage output and to visualize and store acquired data. Signals were sampled at 10 kHz. Liquid junction potentials were not corrected. Patch clamp recordings were guided by a 60x/0.9NA LUMPlanFl/IR Olympus objective. Pipettes and microscope movements were controlled through MP-285 Sutter Instruments. The setup was equipped with a U-RFL-T Olympus fluorescence lamp.

Pyramidal cells were patched in wild type mice since their identification did not require EGFP labeling. Pyramidal cells were identified by their cell shape, localization, input resistance, membrane capacitance and firing pattern. All other cell types were identified based on EGFP expression in reporter mouse lines. Inhibitory and excitatory postsynaptic currents (IPSCs and EPSCs) were recorded in identified cells voltage-clamped at -70 mV. For IPSCs, NBQX (10 μM, Tocris) and CPP (10 μM, Tocris) were added to the bath, and K^+^-based, high Cl^-^ intracellular solution was used (in mM: 127.5 KCl, 11 EGTA, 10 Hepes, 1 CaCl_2_, 2 MgCl_2_, 2 Mg-ATP and 0.3 GTP, pH 7.3 adjusted with KOH). EPSCs were recorded with K^+^-based low Cl^-^ intracellular solution (in mM: 130 K^+^-Gluconate, 10 Hepes, 10 Phosphocreatine-Na, 10 Na-Gluconate, 4 ATP-Mg, 4 NaCl, 0.3 GTP, pH 7.2 adjusted with KOH) and gabazine (10 μM; SR 95531 hydrobromide, Tocris) was bath applied.

Series resistance was continuously monitored in voltage-clamp mode during PSC recordings measuring peak currents in response to small hyperpolarizing pulses. Recordings with series resistance changes of more than 20% were discarded. Series resistances of 35 MOhm were accepted for analyzing PSCs in interneurons and 25 MOhm were accepted for pyramidal cells.

Firing patterns were analyzed in current clamp mode applying 1 s current pulses with 3 s intersweep interval, starting at -200 pA and incrementally increasing the current by 20 pA steps until saturation of action potentials was reached (defined as a decrease in action potential amplitudes). Input resistance was calculated from the steady state voltage step to the first hyperpolarizing current injection for 1 s. Action potential (AP) half width was measured at half amplitude of the AP. Maximal frequency was measured at 1000 pA current injection or directly before saturation was reached. Rheobase was calculated as the minimal injected current that was required to elicit APs in whole-cell mode.

Membrane potential changes upon GRP (Phoenix Pharmaceuticals #027-40, 300 nM if not indicated otherwise) bath application was performed with K^+^-based low Cl^-^ intracellular solution (see above) and with various drug cocktails using the following drugs and concentrations: NBQX (10 μM, Tocris), (R)-CPP (10 μM, Tocris), gabazine (10 μM; SR 95531 hydrobromide, Tocris), CGP55845 (10 μM, Tocris), TTX (1 μM, Tocris), CdCl_2_ (100 μM, Sigma Aldrich) and BW2258U89 (1 μM, Phoenix Pharmaceuticals).

Current clamp recordings were downsampled to 1 kHz for all illustrations and for the analysis of responses to GRP. PSC frequency was calculated in sliding windows of 10 seconds, and baseline frequency was subtracted for each cell. Mean increase in PSC frequency and in membrane potential upon GRP bath application were calculated as difference between mean frequency or membrane potential during baseline and during a 3 min period starting at the time point when GRP reached the bath (based on TAMRA-GRP imaging, see below). A VIP cell was defined as responding to GRP if the mean membrane potential after GRP application was >2 SD above the mean baseline membrane potential and if the membrane potential change was larger than the maximal membrane potential change observed in response to 0 nM GRP. MATLAB was used for offline analysis of all data.

#### HEK 293T live-cell imaging

HEK 293T cells (ATCC Cat# CRL-11268, RRID:CVCL_1926) were transfected with the following plasmids (1.8 ng per well in a 24-well plate): pcDNA3-mRuby2, pAAV-EF1a-FAS-TdTomato, pAAV-CAG-GCAMP6s-WPRE-SV40, pcDNA3.1-mBeRFP. pAAV-CBA-DIO-GCaMP6s-P2A-mBeRFP and pAAV-EF1a-Cre were transfected at a 4:1 ratio.

Cells were imaged 48 hours post transfection. Culture medium was replaced and cells washed three times in imaging buffer (in mM): 125 NaCl, 2 MgCl2, 4.5 KCl, 10 glucose, 20 HEPES pH 7.4. Cells were continuously perfused with imaging buffer via a peristaltic pump. Cells were allowed to equilibrate in the new medium at room temperature until fluorescence had reached a steady state (around 5-10 min). Live-cell imaging was performed at a Leica SP8 X confocal microscope (Harvard NeuroDiscovery Center). During the first imaging session, emission spectra were acquired at constant 473 nm excitation (405 nm excitation to image GCaMP at approximate isosbestic point). Images were taken at 390 to 770 nm emission wavelengths in 10 nm steps with 10 nm imaging bandwidth in a sequence of increasing wavelengths followed by a downward sequence (to balance bleaching effects). To test Ca^2+^ dependence of the fluorescence intensity, 2 mM CaCl_2_ and 10 μM ionomycin (Sigma Aldrich) to promote calcium flux across membranes were added to the imaging buffer. 10 min after, fluorescence was stabilized and a second emission spectrum was acquired as described above.

For quantification of fluorescence changes upon Ca^2+^/ionomycin flow-in, all emission spectra were normalized to the peak fluorescence in Ca^2+^-free buffer. For each well, the area under the curve from 480 to 720 nm was calculated without and with Ca^2+^.

To Compare GCaMP dynamics with and without coexpression of mBeRFP, emission spectra of pAAV-CAG-GCAMP6s-WPRE-SV40- and pAAV-CBA-DIO-GCaMP6s-P2A-mBeRFP-epxressing HEK 293T cells were acquired. Emission spectra were normalized to peak fluorescence at 480-560 nm emission wavelength in Ca^2+^-free buffer and the area under the curve calculated from 480 to 560 nm without and with Ca^2+^.

#### GCaMP-mBeRFP acute slice imaging

For GCaMP imaging upon GRP bath application, auditory or motor cortex of *VIP-IRES-Cre*, *VIP-IRES-Cre;Grpr^wt/y^* or *VIP-IRES-Cre;Grpr^fl/y^* mice were bilaterally injected with 10^12^ GC/ml AAV CBA-DIO-GCAMP-P2A-mBeRFP (400 nl) as described above. To test CRISPR/Cas9 AAVs, AAV CBA-DIO-GCAMP-P2A-mBeRFP was co-injected with AAV pX601-AAV-CMV::NLS-saCas9-NLS-3xHA-bGHpa;U6::BsaI-sgRNA-*Grpr1*, -*Grpr2* or -*lacZ*.

Acute cortical slices were prepared as described for electrophysiological recordings. Slices were imaged with constant fluorescence excitation on an Olympus BX51WI using an Andor Ixon+ camera with Andor Solis Cell A imaging software using a 10x/0.3NA UMPlanFl Olympus objective. The setup was equipped with a U-RFL-T Olympus fluorescence lamp. Excitation filters in the U-MF2 imaging cubes were as follows: 472/30 nm (GCaMP and mBeRFP, Brightline). Emission filter: 520/35 nm (GCaMP), 650/100 nm (mBeRFP broad spectrum), 660/30 nm (reduced bleed through spectrum for mBeRFP: Semrock 3035B modified with FF01-660/30-25 emission filter). Since the GCaMP fluorescence was weak in acute slices at baseline, mBeRFP fluorescence was used to find healthy transfected cells. Each slice was allowed to equilibrate in the imaging chamber for 5 min before video acquisition started. Videos were taken at 32°C.

To verify Ca^2+^ independence of mBeRFP fluorescence in acute brain slices, cell-attached or whole-cell recordings were performed in GCaMP-mBeRFP-expressing VIP cells while videos of GCaMP and mBeRFP fluorescence (472/30 nm excitation and 660/30 nm emission) were acquired. Trains of action potentials were triggered by 10 Hz electrical stimulation for 5 s through the patch pipette. GCaMP and mBeRFP fluorescence were imaged at 5 Hz. Videos were saved as 8-bit TIFs. To image GCaMP at the approximate isosbestic point, a fiber-optic-coupled LED (Thorlabs) was used to excite GCaMP at 405 nm through an optical fiber positioned in the proximity of the imaged cell. For quantifications, mBeRFP and GCaMP (405 and 472/30 nm excitation) mean ΔF/F was calculated during electrical stimulation (5 s duration), and mean mBeRFP and GCaMP (405 nm excitation) ΔF/F were normalized to mean GCaMP (472/30 nm excitation) ΔF/F.

To analyze calcium dynamics upon GRP application, after 3 min of baseline imaging, 300 nM GRP were washed in for 2 min. GRP was washed out for 8 min before 50 mM KCl were bath applied. Video acquisition was stopped after maximum fluorescence was reached.

Video frames were corrected for movements of the slices using custom-written MATLAB software using semi-manual tracking of constant fluorescent markers on the slice.

A custom-written ImageJ script was used to calculate mean fluorescence in manually defined ROIs. Autofluorescence of the slice was calculated from an ROI outside of the injection site, and subtracted from the fluorescence signal. Fluorescence for every ROI was normalized to the peak fluorescence during KCl application to account for variability in GCaMP expression levels per cell. ΔF/F_KCl_ was then calculated by subtracting the average baseline fluorescence.

For quantification of average responses upon GRP application, mean ΔF/F_KCl_ was calculated over 3 min upon GRP bath application, starting with the time point when GRP arrived in the bath (based on TAMRA-GRP fluorescence, see below).

Responding cells were defined as cells with maximal ΔF/F_KCl_ increases upon GRP application of more than 2 SD above maximal ΔF/F_KCl_ during the baseline period. The start of ΔF/F_KCl_ increases after GRP application was defined as the first time point when ΔF/F_KCl_ increased above maximal baseline ΔF/F_KCl_ +2 SD.

#### TAMRA-GRP imaging

TAMRA-GRP was custom-synthetized by Pepscan. TAMRA was attached to the N terminus of GRP. Peptide sequence: Val-Ser-Thr-Gly-Ala-Gly-Gly-Gly-Thr-Val-Leu-Ala-Lys-Met-Tyr-Pro-Arg-Gly-Ser-His-Trp-Ala-Val-Gly-His-Leu-Met-NH2. To test the delay of GRP flow-in on our system, non-fluorescent acute brain slices were imaged with a 650/100 nm emission filter. After 3 min baseline imaging, 300 nM TAMRA-GRP was bath applied for 2 min. The start of TAMRA-GRP arrival in the bath was defined as the first time point when fluorescence increased 2 SD above baseline fluorescence.

#### Sensor engineering and characterization

Sensor engineering and characterization were performed based on previously described protocols (Patriarchi et al., 2018).

The grpLight library was generated using circular polymerase extension cloning (CPEC). The variants were than introduced via PCR for final subcloning into pAAV.hSynapsin1 viral vectors. Active conformations of the sensors were predicted with rosetta_cm protocol of rosetta 3 (version 2015.31). For characterization in cells, HEK 293 cells (ATCC Cat# CRL-1573, RRID:CVCL_0045) were cultured and transfected as in (Patriarchi et al., 2018). Primary hippocampal neurons were freshly isolated and cultured as previously described (Patriarchi et al., 2018). Hippocampal neurons were virally transduced using AAVs (1 x 10^9^ GC/ml) at DIV5, two weeks prior to imaging.

Cells were washed with HBSS (Life Technologies) supplemented with Ca^2+^ (2 mM) and Mg^2+^ (1mM) two times followed by time-lapse imaging with a 40X oil-based objective on an inverted Zeiss Observer LSN710 confocal microscope. For titration curves, apparent affinity (EC50) values were obtained by fitting the data with Hill Equation (Igor). ΔF/F in response to GRP at each concentration was calculated as (*F(t) - F_0_*) / *F_0_* with *F(t)* being the pixel-wise fluorescence value at each time, *t*, and either basal or averaged fluorescence prior to ligand application, *F_0_*. Based on the ΔF/F maps, SNR was calculated as *ΔF/F* x √*F_0_* using a custom-made MATLAB script (Patriarchi et al., 2018).

#### Cannula infusion

Cannulae were surgically implanted to locally infuse GRP into the cortex. Craniotomies were made as described for stereotactic injections. Stainless steel guide cannulae (26 gauge; C315GA/SPC, Plastics One, Roanoke, VA, USA) were positioned in cortical L1 (−2.5 mm anteroposterior, 2 mm mediolateral, −0.2 mm dorsoventral). The gap between the cannula and the skull was filled with a biocompatible transparent silicone adhesive (World Precision Instruments, Kwik-Sil) and allowed to dry for 10 min. The implant was secured with Loctite gel (#454). Hardening of the glue was accelerated by Zip Kicker (Pacer Technology). Dummy cannulae that did not extend beyond the guide cannulae (C315DC/SPC, Plastics One) were inserted to prevent clogging. Mice recovered for 2 weeks. For drug infusions, dummy cannulae were replaced by internal cannula (33 gauge; C315LI/SPC, Plastics One) that extended 1 mm beyond the guide cannulae and were connected to a pump. GRP-TAMRA (2 µl; 3 µg/µl) was infused at 100 nl/min.

#### Fear conditioning

Mice were handled for 7 days, and single-housed for 5-7 days prior to fear conditioning. 2 days before fear conditioning, mice were habituated to the conditioning box for 10 min each day. The behavior boxes consisted of custom-built white plexiglas boxes built around a shock grid floor (Med Associates # ENV-005A). The dimensions of the boxes were 40×30×40cm (WDH). The boxes were equipped with a normal light source (Med Associates # ENV-221CL), a near infrared light source (Med Associates # NIR-200), a speaker (Audax TW025A20) and a camera (Point grey # FL3-U3-13E4M-C and FL3-U3-13Y3M-C). Sound pressure levels were set to 55 ± 1 dBA measured at the bottom of the behavioral box. The shock intensity was set to 0.6 mA consistent with a previous publication (Letzkus et al., 2011). The lights, the shock generator and playback of the sounds were controlled through an Arduino Uno. Sounds were played from an MP3 player (Sparkfun # DEV-12660) and amplified (Sparkfun #BOB-09816).

For fear conditioning, mice were placed into the rectangular box. Lights were switched on 2 min prior to the first sound, to record baseline activity levels. Each mouse was exposed to 15 repetitions of 2 different complex sounds at pseudorandom sequence with inter sound intervals of 60 to 130 sec (in average 82 sec). Each sound consisted of 30 sweeps (either upwards from 5 to 20 Hz or downwards from 20 to 5 Hz). Each sweep was 500 ms long and was followed by a 500 ms sound gap. Each sweep started and ended with a 50 ms ramp to prevent clicking noise of the speakers. Either upsweeps or downsweeps were chosen to serve as the conditioning sound and were paired to a 1 second lasting shock that coincided with the last sweep of the complex sound (conditioned sound, CS^+^). The mice were pseudo-randomly allocated to the behavioral boxes and to the sounds such that the number of control and KO mice receiving either the up or the downsweep as CS^+^ were balanced. Mice were video recorded at 30 frames per second and videos saved as compressed H.264 (AVC) video files.

On the retrieval day, mice were placed into the behavioral boxes equipped with round plastic walls. Light settings were changed to NIR only and the odor was changed to vanilla-odor to reduce contextual fear memory. Sounds were then played either in the same sequence as during the conditioning session or in an inverted sequence to exclude freezing differences inherent to the sequence of the sounds. No foot shocks were applied.

##### Analysis

Freezing was defined as ≥ 2 sec bouts of no movement other than breathing-related movements. Freezing behavior of mice was analyzed with custom-written MATLAB scripts. Freezing was detected based on calculated speed using centroid-based tracking. In brief, the video was thresholded manually, the centroid of the mouse determined for each frame. The speed threshold to detect freezing was semi-automatically determined by playback of video chunks of increasing speed levels until no movement was detected. All putative freezing bouts that were close to the threshold were then automatically played back in MATLAB and manually verified as freezing bouts or excluded. This step was crucial to distinguish between freezing bouts with strong rhythmic body movements due to heavy freezing-associated breathing, and bouts of slight movements including sniffing, certain types of grooming, slight twitching, and ear movements.

Freezing levels and speed during sounds was calculated for the first 29 sec of the sounds only, to exclude the time of the shock.

Discrimination index for the two sounds was calculated as follows: (freezing (CS^+^) - freezing(CS^-^)) / (freezing (CS^+^) + freezing(CS^-^)).

##### Histology

At the end of the experiments, mice were transcardially perfused with PBS and 4% paraformaldehyde. Brains were dissected and sliced on a vibratome (VT1000s, Leica) into 100 μm coronal slices for imaging endogenous fluorescence or 50 μm coronal slices for immunostainings as described above. Mice with spread of AAV-mediated expression into subcortical areas were excluded from the analysis.

#### Photometry

##### Surgery

AAV-CBA-DIO-GCaMP-P2A-mBeRFP or AAVs encoding grpLight were injected as described above into the right hemisphere of the ACx. After removal of the pipette, a syringe needle tip was used to scratch the skull for better adherence of the glue. Tapered fiberoptic cannula implants (MFC_200/230-0.37_2mm_MF1.25_A45) with low autofluorescence epoxy were implanted into the same craniotomy at 0.8-0.95 um depth with the angled (uncoated) side of the fiber tip facing layers 2/3 of the ACx. The gap between the implant and the skull was filled with a biocompatible transparent silicone adhesive (World Precision Instruments, Kwik-Sil) and allowed to dry for 10 min. The implant was secured with Loctite gel (#454). Hardening of the glue was accelerated by Zip Kicker (Pacer Technology). The glue was painted with black nail polish to reduce amount of ambient light collected through the fiber implant.

##### Setup

A 200 μm diameter and 0.37 NA patchcord (MFP_200/220/900-0.37_2m_FCM-MF1.25, low autofluorescence epoxy, Dorics) was used to connect fiber implants to a Dorics filter cube with built-in photodetectors for blue excitation light (465-480 nm) green (500-540 nm) and red (580-680 nm) emission light (FMC5_E1(465-480)_F1(500-540)_E2(555-570)_F2(580-680)_S, Dorics). Signals from the photodetectors were amplified with Dorics amplifiers with a gain of 1-10x in DC mode and acquired using a Labjack (T7). LJM Library (2018 release) was installed to allow communication between MATLAB and Labjack through a USB connection. The voltage output from the LED drivers was amplitude modulated at 171 (470 nm excitation of GCaMP, mBeRFP and the GRP sensor) and 228 (565 nm excitation of TAMRA-GRP) Hz to filter out ambient light and bleed-through emission. Amplitude modulation was programmed in MATLAB. 470 nm LEDs (M470F3, Thorlabs; LED driver LEDD1B, Thorlabs) and 565 nm LEDs (M565F3, Thorlabs, LED driver LEDD1B, Thorlabs) were used. Light power at the patchcord tip was set to oscillate between 38 and 75 μW for 470 nm excitation and 23-38 μW for 565 nm excitation (min and max of the amplitude modulation sine wave).

For synchronization with behavior data, timestamps of each collected video frame as well as signals from the Arduino channels for light, shock and sound were collected with the Labjack synchronously.

##### Recordings

Mice were handled to fiber attachment for 7 days prior to behavioral training. Mice were allowed to move freely on the hand while cleaning the fiber implant with ethanol and while connecting the patchcord to the fiber implant. mBeRFP fluorescence was used to verify proper connection and stable fluorescence across days. We found that GCaMP and mBeRFP expression were stable after 4 weeks of expression, and thus all data were collected after at least 4 weeks of expression. Photometry data were sampled at 2052 Hz for all channels, and saved in 1 sec chunks. Collection started at least 10 secs before connection to determine autofluorescence from the patchcord. Since bleaching of GCaMP and mBeRFP were strongest during the first 2 days of recordings, photometry data from the 2 habituation sessions prior to fear conditioning were not used. On the following days, the first 10-30 sec of recordings after patchcord connection were discarded to exclude the initial drop in fluorescence due to bleaching and due to handling of the mouse each day. Traces with sudden drops in mBeRFP signal during the behavioral session pointed to detachment or coiling of the fiber and were discarded. To reduce bleaching during the behavioral settings, LEDs were switched off during sounds 6 to 10. Thereby we were able to achieve constant fluorescence levels on conditioning and retrieval day.

##### Analysis

Since the collected GCaMP fluorescence but not ambient light was amplitude-modulated by a 171 Hz sine wave, we analyzed GCaMP fluorescence by calculating the power at 171 ± 7.5 Hz using online and offline analysis with custom-written MATLAB scripts based on a previous publication (Owen and Kreitzer, 2019). Power of the fluorescence signals at 171 ± 7.5 Hz was calculated over a 200 sample point sliding window with 180 sample point overlap. The resulting trace was corrected for bleaching: based on recordings from 3 mice without fluorophores, we found that the majority of bleaching during the behavioral sessions could be ascribed to autofluorescence bleaching. Therefore, we calculated an exponential fit for the minima of GCaMP and mBeRFP fluorescence traces (minima based on a sliding window of 5000 data points), corrected for the exponential drop and subtracted autofluorescence. We normalized GCaMP fluorescence to the mean expression level of mBeRFP (Fig 6). For Fig. S6, GCaMP and mBeRFP fluorescence were instead normalized to minimal fluorescence levels at the end of the conditioning session to compare GCaMP and mBeRFP fluorescence and exclude movement artifacts.

#### Statistics

Normally distributed data with equal variance were compared using two-sample or paired t-tests, and data were shown as mean ± SEM. Normally distributed data with unequal variance were compared using the two-sample t-test for unequal variance and data were shown as mean ± SEM. Non-normally distributed data were compared using the Mann-Whitney-U test and data were shown as median (IQR). The Shapiro-Wilk test was used to test for normal distribution of data. Normally distributed data were tested for homogeneity of variance using the F-test. P-values were corrected for familywise error rates with the Holm-Bonferroni test where applicable.

Behavioral data were compared using two-way ANOVA and Tukey’s posthoc test if main interactions were significant.

Cumulative frequency distributions were compared using two-sample Kolmogorov-Smirnov tests. Graphs were made with custom-written scripts in MATLAB. The figures were assembled in Illustrator (Adobe). The following code was used for p-values in the figures: *<0.05; **<0.01; ***<0.001.

## SUPPLEMENTAL INFORMATION

Document S1. Figures S1-S7.

---

**Figure S1:**
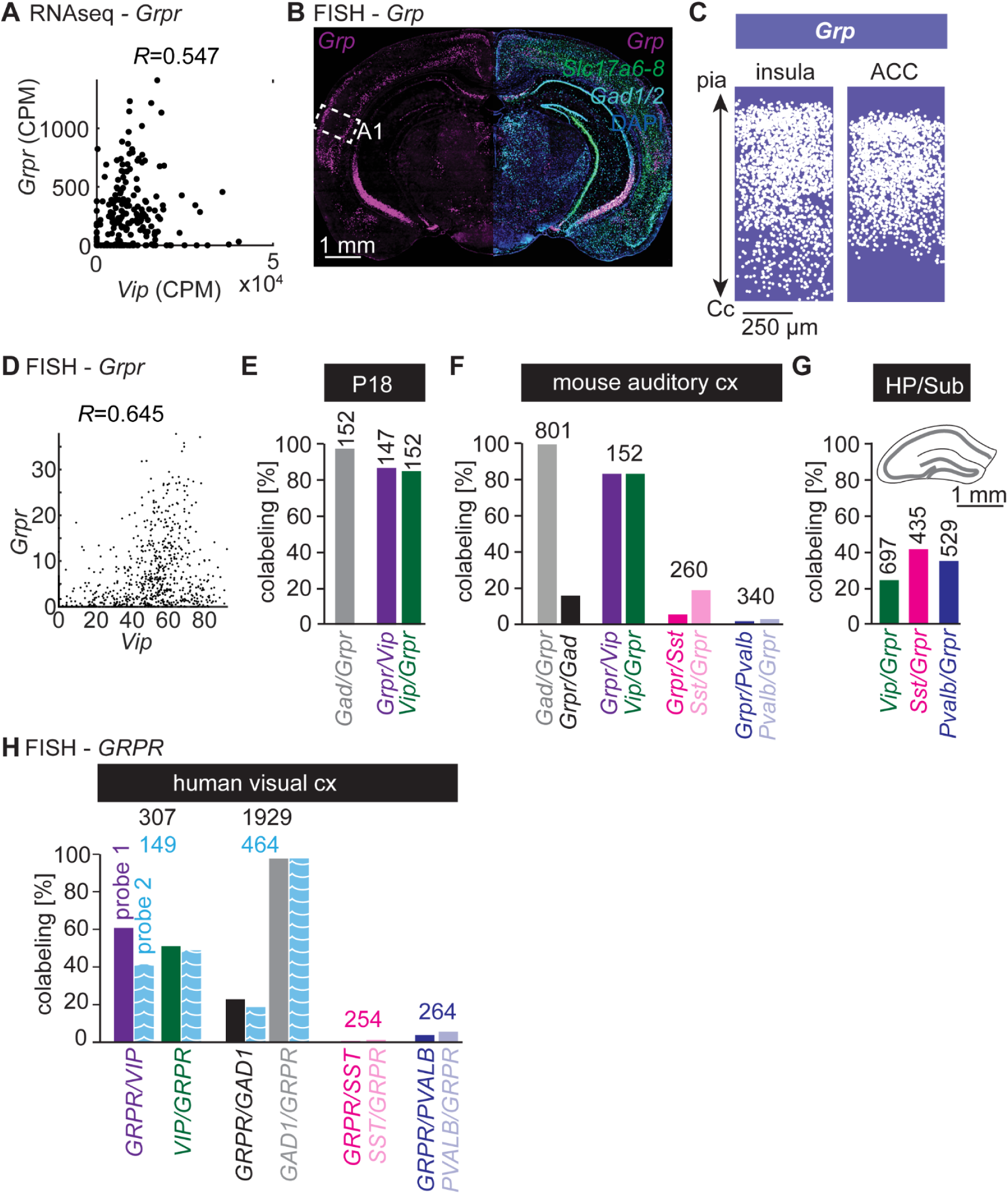
**Cortex-wide cell type-specific expression of GRP and its receptor in mice and humans. Related to Figure 1.** **A**, Correlation of *Grpr* and *Vip* expression levels (counts per million, CPM) in the mouse visual cortex based on RNA sequencing data published in Tasic *et al*. 2016. **B**, Representative epifluorescent image of auditory cortex labeled for *Grp*, glutamatergic markers (*Slc17a6-8*) and GABAergic markers (*Gad1,2*) using FISH. **C**, Overlay showing the locations of all identified *Grp^+^* cells in anterior insula and anterior cingulate cortex (ACC). N = 8 (insula) and 5 (ACC) slices from 4-7 hemispheres per area. **D**, Correlation of *Grpr* and *Vip* expression levels (percent coverage per cell body) across all examined cortical areas. n = 916 cells. Same data set as in Figure 1D. **E**, Quantification of *Grpr* coexpression with indicated genes in auditory cortex of mice at postnatal day 18. The numbers of counted cells per brain area are indicated above bars. N = 3 mice each. **F**, Quantification of *Grpr* coexpression with indicated genes in auditory cortex of adult mice. The numbers of counted cells per brain area are indicated above bars. N = 3 mice each. **G**, Quantification of *Grpr* coexpression with indicated genes in the hippocampus/subiculum of adult mice. The numbers of counted cells per brain area are indicated above bars. N = 3 mice each. **H**, Quantification of *GRPR* coexpression with indicated markers in human visual cortex. The numbers of counted cells per brain area are indicated above bars. Blue patterned bars and blue numbers: quantification based on FISH with a second *GRPR* probe targeting a distinct *GRPR* sequence.

**Figure S2.1:**
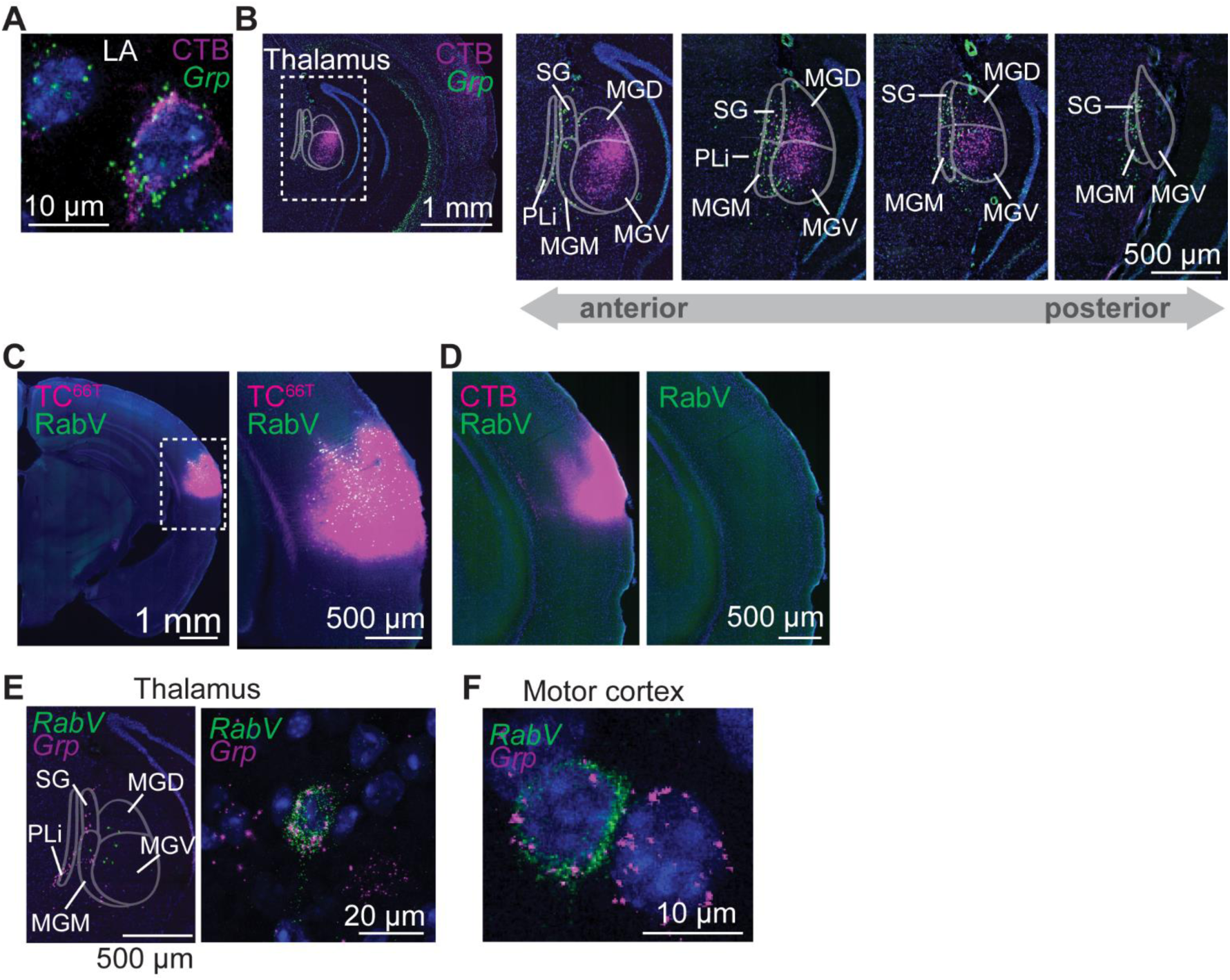
**Putative local and long-range sources of synaptic and extrasynaptic GRP. Related to Figure 2.** **A**, Confocal image of a retrogradely labeled and *Grp*-positive cell in the lateral amygdala (LA) after CTB injection into the auditory cortex. **B**, Representative epifluorescent images of retrogradely labeled cells in several thalamic nuclei along the anterior-posterior axis and FISH against *Grp* after CTB injection into the auditory cortex. **C**, Representative epifluorescent image of SADΔG-EnVA-H2B-EGFP (RabV) and CTB after injection of helper viruses AAV DIO-TC^66T^-mCherry and AAV DIO-oG, followed by injection of RabV and CTB into auditory cortex of C57BL/6J mice. **D**, Representative epifluorescent image of SADΔG-EnVA-H2B-EGFP (RabV) and TC^66T^-mCherry (TC^66T^) expression after injection of helper viruses AAV DO-TC^66T^-mCherry and AAV DIO-oG, followed by injection of RabV into auditory cortex of C57BL/6J mice. Right: magnification of the indicated area. **E**, Representative confocal images of FISH against *RabV-gp1 (RabV)* and *Grp* in auditory thalamus after injection of AAV DIO-TC^66T^-mCherry and AAV DIO-oG, followed by injection of SADΔG-EnVA-H2B-EGFP into auditory cortex of *Vip-IRES-Cre* mice. **F**, Representative confocal image of FISH against *RabV-gp1 (RabV)* and *Grp* in motor cortex after injection of AAV DIO-TC^66T^-mCherry and AAV DIO-oG, followed by injection of SADΔG-EnVA-H2B-EGFP into motor cortex of *Vip-IRES-Cre* mice.

**Figure S2.2.**
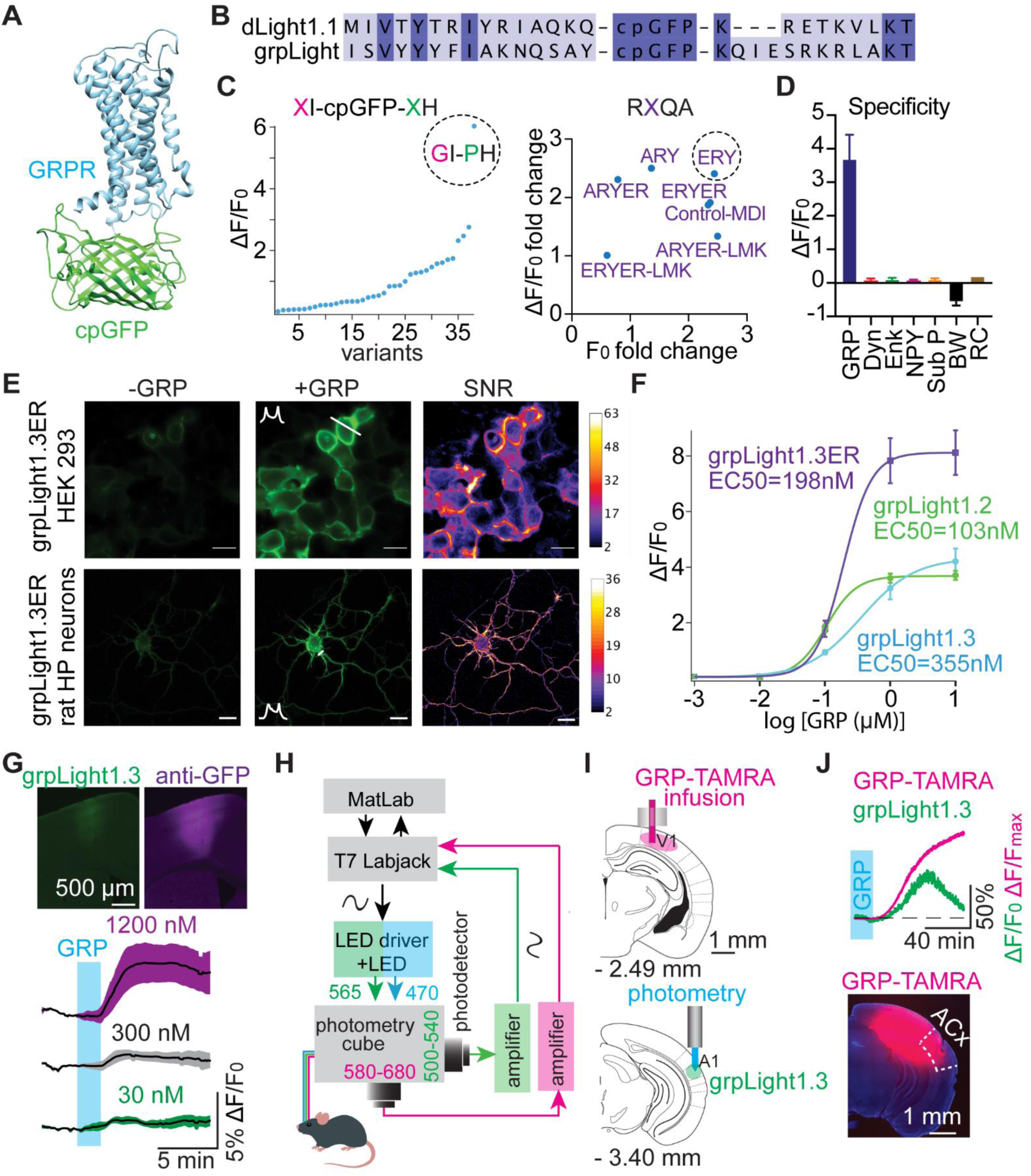
**A**, Simulated structure of grpLight1 consisting of GRPR and cpGFP module. **B**, Sequence alignment of dLight1.1 and grpLight1 adjacent to intracellular loop 3 linker **C**, Screening results for variants in linker between cpGFP and GRPR (*left*) and in intracellular loop 2 (*right*). *Left,* Fluorescence changes to 1 µM GRP. *Right*, Fold change of basal fluorescence and fluorescence changes compared to control grpLight prior to and following application of 1 µM GRP. **D**, Pharmacological specificity of grpLight1.2. Fluorescence changes (ΔF/F_0_) to GRP (100 nM), dynorphin (DYN, 100 µM), enkephalin (ENK, 100 µM), neuropeptide Y (NPY, 100 µM), substance P (Sub P, 100 µM), GRPR antagonist BW2258U89 (500 µM) and GRPR antagonist RC3095 (500 µM). **E**, Expression of grpLight1.3ER in HEK 293T cells and rat hippocampal neurons. Fluorescence response following application of 10 µM GRP and signal-to-noise ratio are shown. Scale bar, 10 µm. **F**, *In situ* titration of GRP on rat hippocampal neurons expressing grpLight variants 1.2, 1.3 and 1.3ER. Half maximal affinities (EC50) are shown. Data were fitted with Hill Equation. Mean ± SEM. **G**, Representative epifluorescent images of grpLight1.3 expression in mouse motor cortex with and without immunostaining (*top*). Fluorescence changes of grpLight1.3 in acute motor cortex slices following bath application of indicated concentrations of GRP. n= 3, 7 and 7 slices (from top to bottom). Mean ± SEM. **H**, Schematic drawing of photometric setup for dual-color fiber photometry. **I**, Schematic of cannula placement in visual cortex and fiber photometry in auditory cortex. **J**, *Top*, Analysis of TAMRA-GRP and grpLight1.3 fluorescence changes recorded with dual-color fiber photometry, following cannula infusion of 6 µg TAMRA-GRP (2 µl at 100 nl/min, blue bar) into visual cortex as shown in J. *Bottom*, Epifluorescent image of GRP-TAMRA fluorescence in auditory cortex (ACx) surrounding the fiber track 1 hour after cannula infusion.

**Figure S3:**
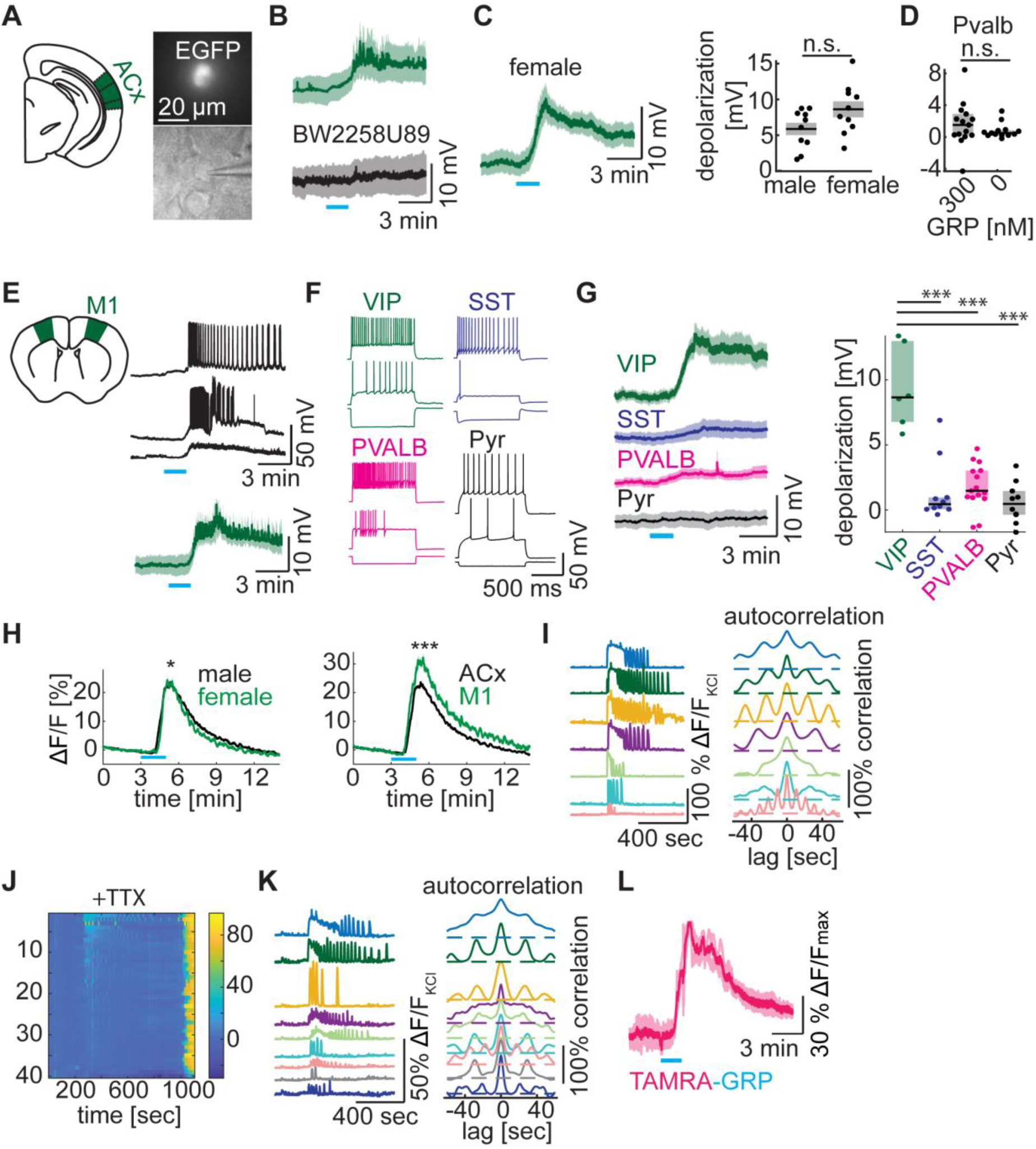
**GRP depolarizes cortical VIP cells and induces calcium signaling. Related to Figure 3.** **A**, Epifluorescent and Dodt gradient contrast images of a patched EGFP^+^ cell in an acute auditory cortex slice. **B**, Average depolarization of auditory cortex VIP neurons upon application of 300 nM GRP with or without 10 µM GRPR antagonist BW2258U89. Mean ± SEM across 10 (*top*) and 6 (*bottom*) cells from male mice. Bath contains NBQX, CPP, gabazine and CGP. **C**, *Left*, Average depolarization of auditory cortex VIP neurons upon application of 300 nM GRP. Mean ± SEM across 10 cells from female mice. Bath contains NBQX, CPP, gabazine, CGP and TTX. *Right*: Amplitude of membrane potential changes in VIP cells of male and female mice. Mean ± SEM. n = 10 cells each. 2-sample t-test: t(18)=1.95, p=0.07. **D**, Quantification of membrane potential changes of PVALB cells in the auditory cortex following bath application of indicated concentrations of GRP (median and IQR). Bath contains NBQX, CPP, gabazine, CGP and TTX. n = 15 and 13 cells. Mann-Whitney U test: U=76, p=0.33. **E**, Exemplary (*top*) and average (*bottom*) depolarization of VIP cells following application of 300 nM GRP in acute motor cortex slices from male mice. Mean ± SEM across 10 cells. Bath contains NBQX, CPP, gabazine, CGP. **F**, Representative firing patterns of motor cortex VIP, SST, PVALB and pyramidal (Pyr) cells as indicated upon -200 pA current injection (*bottom*), at AP threshold (*middle*), and at maximal firing frequency (*top*). **G**, Average time course (*left*) and amplitude (*right*) of the membrane potential changes in the indicated cell types in motor cortex following GRP application. Mean ± SEM (left) and median/IQR (right) across 6 VIP, 10 SST, 15 PVALB and 10 pyramidal cells. Bath contains NBQX, CPP, gabazine, CGP and TTX. Comparison of depolarizations across cell types (Mann-Whitney U test): SST: U = 2, p = 0.001; PVALB: U = 0, p = 0.001; pyramidal: U = 0, p = 0.001. **H,** Average GCaMP fluorescence changes (normalized to KCl response) across all recorded VIP cells in acute cortex slices of male and female auditory cortex (*left*) and male auditory and motor cortex (*right*). Mean ± SEM. Mann-Whitney U test: male vs. female: U = 413360, p = 0.047; n = 1510 and 580 cells in 37 and 14 slices respectively; auditory vs. motor cortex: U = 233659, p < 0.0001, n = 405 cells in 8 motor cortex slices. **I**, GCaMP fluorescence changes (normalized to KCl response, *left*) in 7 exemplary VIP cells responding to GRP application with oscillatory Ca^2+^ dynamics. Autocorrelation (*right*) showing different ‘oscillation’ frequencies. Same dataset as in Figure 3H. **J**, Heatmap of fluorescence changes (expressed relative to KCl fluorescence) across all imaged VIP cells in an exemplary acute auditory cortex slice. Bath contains NBQX, CPP, gabazine, CGP and TTX. **K**, GCaMP fluorescence changes (normalized to KCl response, *left*) and autocorrelation (*right*) of 9 exemplary VIP cells responding to GRP application with oscillatory Ca^2+^ dynamics. Same dataset as in J. **L**, Average TAMRA-GRP fluorescence changes upon bath application of fluorescently tagged GRP (TAMRA-GRP). Mean ± SEM across 3 slices.

**Figure S4:**
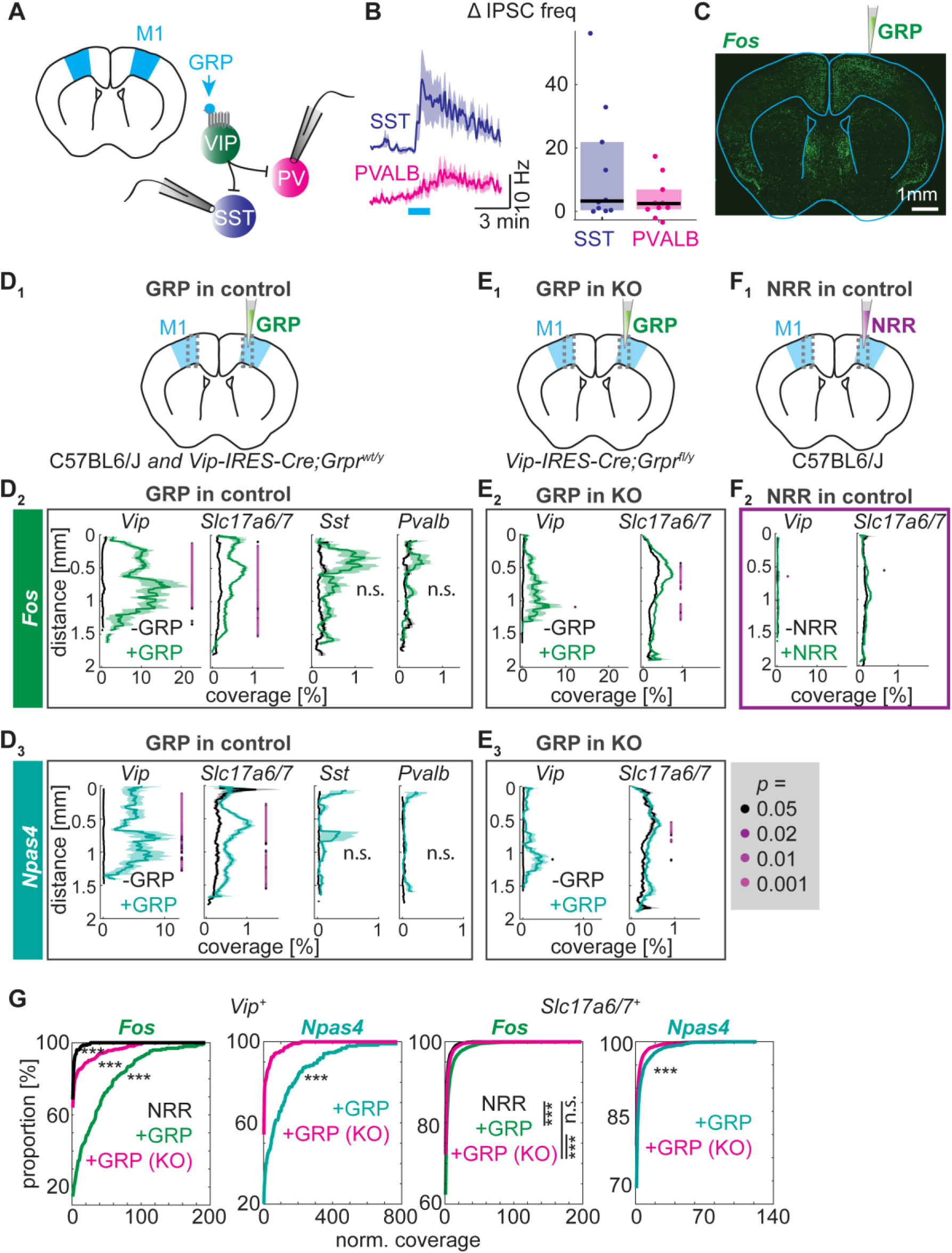
**GRP disinhibits the cortex and induces immediate early gene expression. Related to Figure 4.** **A**, Schematic of voltage-clamped motor cortex SST and PVALB cells and bath application of GRP. **B**, Average time course (*left*, mean ± SEM) and magnitude (*right,* median and IQR) of IPSC frequency changes upon GRP bath application. n = 10 cells per group. Bath contains NBQX and CPP. **C**, Representative epifuorescent image of FISH against *Fos* after unilateral injection of 3 µM GRP, as schematized, into the right motor cortex in anaesthetized mice. **D-F**, *Fos* (**D_2_**, **E_2_**, **F_2_**) and *Npas4* (**D_3_**, **E_3_**, **F_3_**) expression levels (defined as cell area covered with FISH labeling) were quantified in *Vip^+^, Sst^+^, Pvalb^+^* and glutamatergic cells across all cortical layers for the right (green/turquoise) and left (black) motor cortices (mean ± SEM). 3 µM GRP or NRR were injected into the right motor cortex of anaesthetized control or conditional *Grpr* knockout mice as indicated in the schematics (**D_1_**, **E_1_**, **F_1_**). n=2-3 slices from 3-5 mice per group. Statistics: bonferroni-corrected p-values for comparison of mean expression in 150 µm bins. P-values shown as color coded dots. **D_2_**, N = 398 (*Vip*), 15108 (*Slc17a6,7*), 921 (*Sst*) and 1300 (*Pvalb*) cells. **E_2_**, N = 673 (*Vip*) and 13477 (*Slc17a6,7*) cells. **F_2_**, N = 455 (*Vip*) and 13217 (*Slc17a6,7*) cells. **D_3_**, N = 417 (*Vip*), 10501 (*Slc17a6,7*), 677 (*Sst*) and 748 (*Pvalb*). **E_3_**, 451 (*Vip*) and 6371 (*Slc17a6,7*) cells. **G**, Cumulative distribution of expression levels (percent coverage) based on data shown in **D**-**F**. Expression levels in the right hemisphere were normalized to the mean expression levels measured in the left hemisphere for each slice to account for differences in baseline expression levels. Comparisons: Two-sample Kolmogorov-Smirnov test.

**Figure S5:**
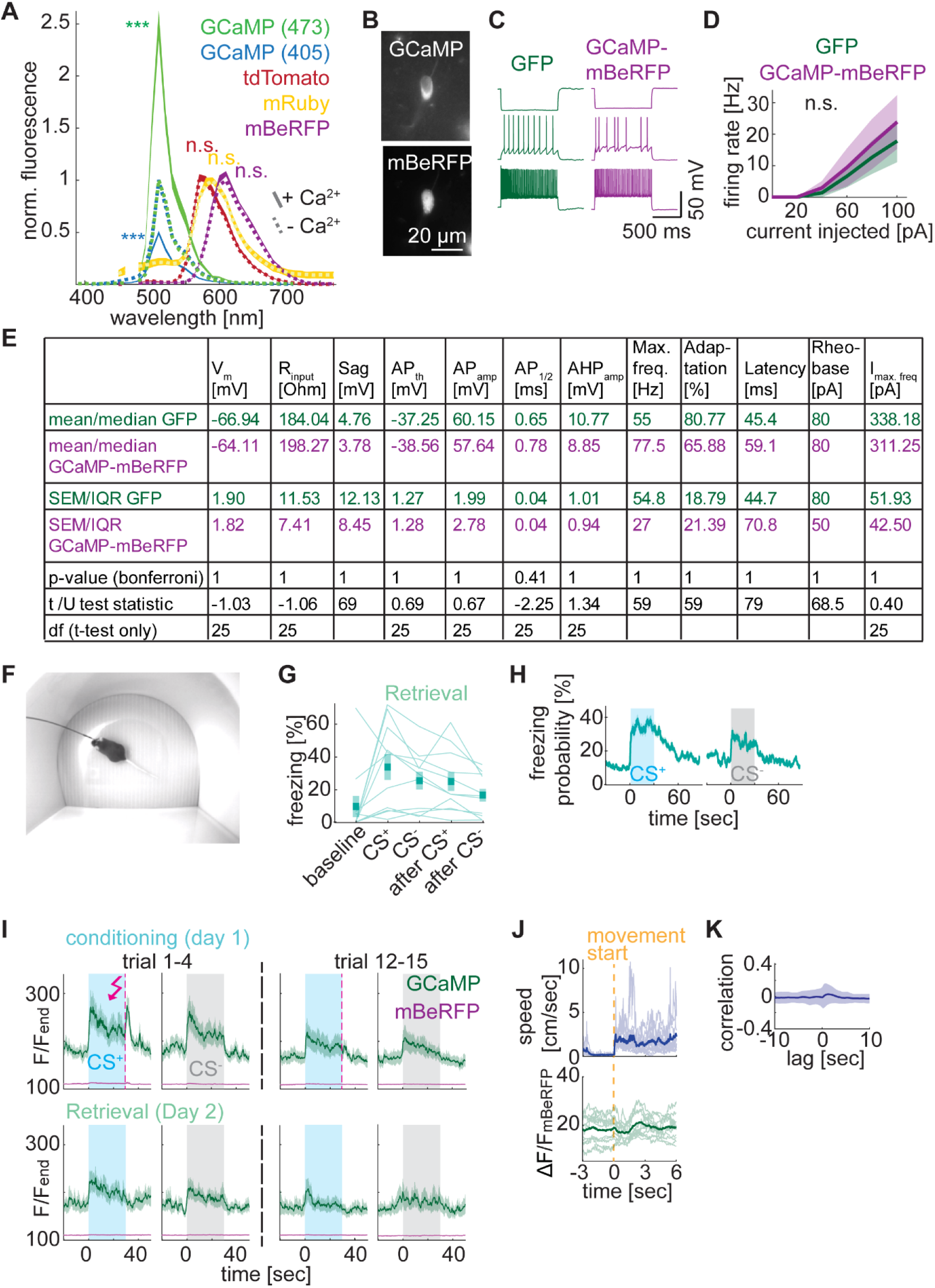
**ACx VIP cells encode novel sounds and shocks during fear conditioning. Related to Figure 5.** **A**, Fluorescence emission spectrum of GCaMP, mBeRFP, tdTomato and mRuby expressed in HEK 293T and measured with (mean ± SEM) or without (dashed line) application of Ca^2+^/ionomycin. Fluorescence was normalized to the maximum fluorescence in 0 Ca^2+^. N = 6 wells each. Excitation = 473 nm (405 nm where indicated). Fluorescence of Ca^2+^-bound GCaMP (473) is shown 10x smaller for better visualization. Comparison of Ca^2+^-dependent and -independent fluorescence: Bonferroni-corrected paired t-tests: GCaMP (473): t(5) = 15.15, p = 0.0001; GCaMP (405): t(5) = -42.51, p < 0.0001; mBeRFP: t(5) = 2.54, p = 0.16; mRuby: t(5) = 0.29, p = 0.78; tdTomato: t(5) = 0.94, p = 0.78. **B**, Representative epifluorescent images of an auditory cortex VIP cell expressing GCaMP-P2A-mBeRFP. **C**, Representative firing patterns of VIP cells expressing GFP or GCaMP-P2A-mBeRFP as indicated upon -200 pA current injection (*bottom*), at AP threshold (*middle*), and upon 100 pA current injection (*top*). **D**, Firing rates of VIP cells expressing GFP or GCaMP-P2A-mBeRFP upon injection of increasing current steps. Mean ± SEM. n = 11 GFP-, 16 GCaMP-P2A-mBeRFP-expressing VIP cells. n-way ANOVA, main effect for GFP vs. GCaMP-P2A-mBeRFP: p = 0.26, F = 1.28. **E**, Active and intrinsic electrophysiological properties of GFP- (n = 11) and GCaMP-P2A-mBeRFP-expressing (n = 16) cortical VIP cells. Statistics: 2-sample t-test and Mann-Whitney U test; Bonferroni-corrected p-values. **F**, Exemplary video frame of an implanted *Vip-IRES-Cre* mouse during fear retrieval session and fiber photometric recording. **G**, Auditory fear memory retrieval in implanted *Vip-IRES-Cre* mice used for photometric recordings, measured as the percentage of time spent freezing averaged across 15 presentations of CS^+^ and CS^-^ on the retrieval day. Mean ± SEM. N = 11 mice. **H**, Time course of average freezing probability across all CS^+^ and CS^-^ during fear memory retrieval. Mean ± SEM. Same mice as in G. **I**, GCaMP and mBeRFP fluorescence (normalized to baseline fluorescence at the end of the conditioning session after subtraction of autofluorescence) measured around presentation of conditioned (CS^+^, blue) and unconditioned sounds (CS^-^, grey) and shocks (dashed pink lines) early (trial 1-4) and late (trial 12-15) on the conditioning (top) or retrieval (bottom) day. Mean ± SEM across 11 mice. Same dataset as in Figure 5I. **J**, Speed (*top*) and GCaMP fluorescence changes (*bottom*, normalized to mBeRFP fluorescence) aligned to movement initiation after periods of spontaneous freezing (in the absence of sounds) during trials 1-4 of the retrieval session across 11 mice (average in bold; traces for each mouse in bright colors). **K**, Cross-correlation of speed and GCaMP fluorescence changes (normalized to mBeRFP fluorescence). Mean ± SEM across 11 mice.

**Figure S6:**
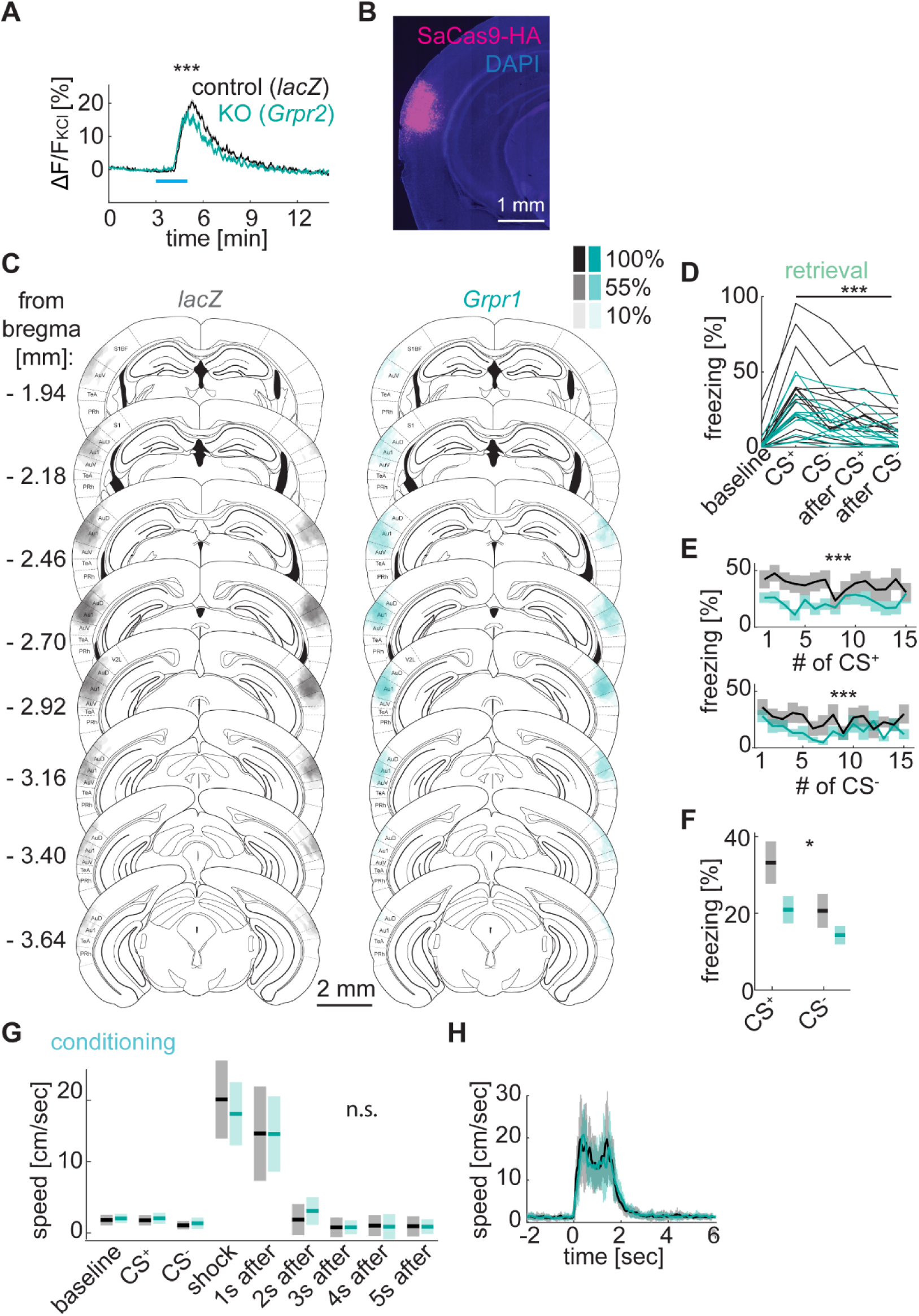
**GRP-GRPR signaling in the ACx enhances fear memories. Related to Figure 6.** **A**, GCaMP fluorescence changes measured in acute brain slices upon GRP application for VIP cells infected with AAV encoding saCas9-sgRNA targeting either *Grpr* (sgRNA-*Grpr2*) or *lacZ* (control) (fluorescence is normalized to KCl fluorescence, mean ± SEM). Mann-Whitney U test: U = 80817, p < 0.0001, n = 602 and 333 cells in 10 and 9 slices for *lacZ* and *Grpr2,* respectively. **B**, Representative epifluorescent image of an immunostaining against saCas9-HA after injection of AAV CMV-saCas9-HA-U6-sgRNA into auditory cortex. **C**, Quantification of bilateral saCas9-HA expression in mice with focal AAV injection into auditory cortex encoding sgRNA targeting *lacZ* (control, grey) or *Grpr* (KO, turquoise). Color-code shows percentage of mice with saCas9-HA expression. **D**, Auditory fear memory retrieval, measured as the percentage of time spent freezing averaged across 15 presentations of CS^+^ and CS^-^ on the retrieval day. Same dataset as in Figure 6E. **E** Auditory fear memory retrieval, measured as the percentage of time spent freezing across 15 CS^+^ and CS^-^. Mean ± SEM. 2-way ANOVA for CS^+^: p < 0.0001, F = 30.9 (genotype) and p = 0.85, F = 0.61 (stimulus number), no significant interaction of genotype and stimulus number; CS^-^: p = 0.0001, F = 15.71 (genotype) and p = 0.20, F = 1.31 (stimulus number), no significant interaction of genotype and stimulus number. **F**, Auditory fear memory retrieval, measured as the percentage of time spent freezing averaged across 15 presentations of CS^+^ and CS^-^ on the retrieval day. Baseline freezing level was subtracted. Mean ± SEM. 2- way ANOVA: main effect of genotype: p = 0.029, F = 5.05, no significant interaction of genotype and stimulus. **G**, Average locomotion during baseline, first CS^+^, CS^-^ and shock, and following the first shock. Mean ± SD across 15 mice per group. 2-way ANOVA: main effect of genotype: p = 0.93, F = 0.01, no significant interaction between genotype and condition. **H**, Average locomotion before, during and after the first 4 foot shocks. Mean ± SD across 15 mice per group.

**Figure S7:**
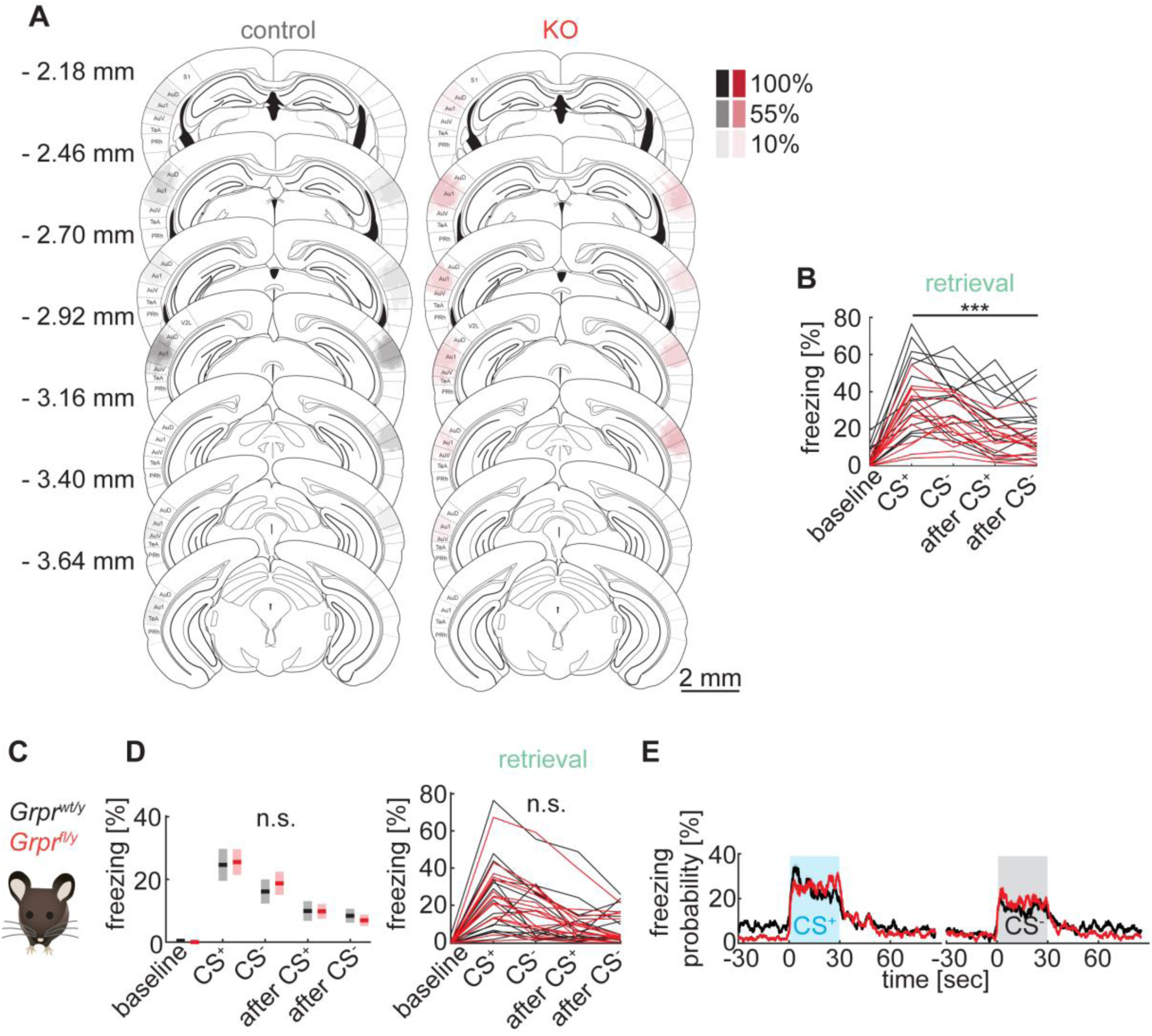
**Impaired fear memory in mice with conditional KO of GRPR in the auditory cortex. Related to Figure 7.** **A**, Quantification of bilateral Cre-mCherry expression in *Grpr^wt/y^* (control) or *Grpr^fl/y^* (KO) mice with focal AAV injection into auditory cortex encoding hSyn-Cre-mCherry. Only the main injection sites were analyzed. Color-code shows percentage of mice with Cre-mCherry expression hotspot. **B**, Auditory fear memory retrieval, measured as the percentage of time spent freezing averaged across 15 presentations of CS^+^ and CS^-^ on the retrieval day. Same dataset as in Figure 7C. **C**, Fear memory was examined in uninjected *Grpr^wt/y^* and *Grpr^fl/y^* mice. **D**, Auditory fear memory retrieval, measured as the percentage of time spent freezing averaged across 15 presentations of CS^+^ and CS^-^ on the retrieval day. 2-way ANOVA: main effect of genotype: p=0.84, F=0.04, no significant interaction between genotype and stimulus; n=16 mice per group. **E**, Timecourse of average freezing probability across all CS^+^ and CS^-^ presentations during fear memory retrieval. Mean ± SEM.

## Notes

### Competing Interest Statement

The authors have declared no competing interest.

